# Superinfection with intact HIV-1 results in conditional replication of defective proviruses and nonsuppressible viremia in people living with HIV-1

**DOI:** 10.1101/2025.04.04.647291

**Authors:** Vivek Hariharan, Jennifer A. White, Filippo Dragoni, Emily J. Fray, Nicholas Pathoulas, Milica Moskovljevic, Hao Zhang, Anushka Singhal, Jun Lai, Subul A. Beg, Eileen P. Scully, Elizabeth A. Gilliams, David S. Block, Jeanne Keruly, Richard D. Moore, Janet D. Siliciano, Francesco R. Simonetti, Robert F. Siliciano

## Abstract

During replication of some RNA viruses, defective particles can spontaneously arise and interfere with wild-type (WT) virus replication. Recently, engineered versions of these defective interfering particles (DIPs) have been proposed as an HIV-1 therapeutic. However, DIPs have yet to be reported in people with HIV-1 (PWH). Here, we find DIPs in PWH who have a rare, polyclonal form of non-suppressible viremia (NSV). While antiretroviral therapy (ART) rapidly reduces viremia to undetectable levels, some individuals experience sustained viremia due to virus production from cell clones harboring intact or defective proviruses. We characterized the source of NSV in two PWH who never reached undetectable viral load despite ART adherence. Remarkably, in each participant, we found a diverse set of defective viral genomes all sharing the same fatal deletions. We found that this paradoxical accumulation of mutations by viruses with fatal defects was driven by superinfection with intact viruses, resulting in mobilization of defective genomes and accumulation of additional mutations during untreated infection. We show that these defective proviruses interfere with WT virus replication, conditionally replicate, and, in one case, have an R_0_ > 1, enabling *in vivo* spread. Despite this, clinical outcomes show no evidence of a beneficial effect of these DIPs.

## Introduction

Antiretroviral therapy (ART) halts HIV-1 replication, reducing plasma HIV-1 RNA to below the limit of detection in most people with HIV-1 (PWH)^1–3^. However, ART is not curative due to the persistence of HIV-1 in a latent form in long-lived resting memory CD4^+^ T cells^4–8^. This latent reservoir carrying infectious, replication-competent proviruses can cause viral rebound when ART is interrupted^9,10^. The reservoir decays very slowly (t_1/2_ = 3.7 years)^11,12^, necessitating lifelong ART. After many years on ART, the frequency of CD4^+^ T cells with inducible, replication-competent virus may actually begin to increase^13^, likely due to antigen-driven proliferation^14–16^.

In most PWH on ART, more than 90% of proviruses are defective, principally due to large deletions and/or APOBEC3G/F-mediated G-to-A hypermutation^17–19^. Defective proviruses are generated throughout the course of untreated infection^20^. Missense mutations can arise due to the low fidelity of HIV-1 reverse transcriptase^21^. Some of these mutations are fatal, but others allow rapid evolution and escape from immune responses^22–26^. Proviruses with large internal deletions are common in PWH on ART. They result from the high propensity of HIV-1 to undergo recombination during reverse transcription through template switching events between the two viral genomes packaged in each virion^27^. Aberrant switching events commonly give rise to internal deletions^17,18,20^. Another major class of defective proviruses are those with G-to-A hypermutation introduced by APOBEC3G/F^17,18,28–30^. Hypermutation introduces multiple stop codons into most ORFs, causing loss of replication competence^18,30^. Proviruses with fatal defects are an evolutionary dead end; therefore, their sequences are unique and should not contribute to further evolution of intra-host viral lineages. However, they can persist and accumulate due to cell proliferation, resulting in cells with identical defective proviral sequences sharing the same integration site^31–36^.

Some defective proviruses are unable to produce viral gene products due to internal deletions, stop codons, and APOBEC3-induced mutations which affect the transcription and translation of viral genes. However, a fraction of defective proviruses can express HIV-1 RNA, and some can produce viral proteins and release virions^36–39^. We have recently shown that defective proviruses with deletions or point mutations in the 5’-leader region can cause non-suppressible viremia (NSV), a clinical phenomenon in which some PWH experience persistent low-level viremia^36^. NSV is not due to suboptimal adherence to ART or drug resistance leading to viral replication, but rather due to the release of virions from clonally expanded CD4^+^ T cells carrying infectious or defective proviruses^33,36,40–42^. The detection of low-level viremia on ART can also be an indication of virological failure, and thus understanding its underlying cause is both a research and clinical priority^43^.

Engineered defective viral genomes have been proposed as an anti-viral strategy^44–46^ because these viruses can impact the replication of wildtype (WT) virus through an interference phenomenon originally described by von Magnus for the influenza virus^47^. These interfering viruses, termed defective interfering particles (DIPs), are incapable of self-replication, but can specifically interfere with the replication of non-defective homologous virus in co-infected cells^48–50^, thereby reducing cytotoxicity, viremia, and pathogenesis. Defective interfering genomes are defined by four biological properties: (i) they retain some genes encoding for structural proteins, (ii) they contain a portion of the viral genome, (iii) they can reproduce conditionally only with the WT virus and, (iv) they interfere with the replication of non-defective homologous virus^51^. DIPs have been observed to arise spontaneously for numerous RNA virus infections^52–55^. Despite the generation of defective proviruses during HIV-1 infection and extensive sequencing of HIV-1, HIV-1 DIPs have not been reported in PWH^46^. However, recent work from Pitchai, Tanner, and colleagues has shown that *in vitro* engineered versions of HIV-1 DIPs, therapeutic interfering particles (TIPs), can act as an antiviral therapeutic by reducing the HIV-1 viral load in humanized mice and non-human primate models^50^.

Here, we demonstrate the presence of DIPs in PWH who have an unusual polyclonal form of NSV that prevents suppression of viremia to undetectable levels. We describe how DIPs contribute to viremia and consider the clinical implications.

## Results

### Persistent viremia despite long-term adherence to effective ART

The participants in this study were referred for evaluation due to sustained detectable viremia despite optimal adherence to ART.

Participant 1 (P1) is a Caucasian male diagnosed with HIV-1 in late 2019, presenting with plasma HIV-1 RNA of 1,240,000 copies/mL and a CD4^+^ T cell count of 8 cells/µL. He was promptly treated with bictegravir, emtricitabine, and tenofovir alafenamide resulting in a 3-log reduction in HIV-1 RNA within 3 months (Fig. 1a). However, despite a significant increase in CD4^+^ T cell count to 132 cells/μL, plasma HIV-1 RNA remained between 100 and 1,000 copies/mL for more than three years. Viral and proviral genotyping revealed only a G190A mutation in *pol*, which did not confer resistance to his treatment regimen. The treatment regimen was switched to emtricitabine, tenofovir alafenamide, darunavir (with cobicistat), plus dolutegravir (50 mg daily). Despite the change in treatment, HIV-1 RNA copies remained stable (100-500 copies/mL). Measurement of drug concentrations showed low trough levels for dolutegravir, prompting an increase to 50 mg dolutegravir twice daily. One month later, dolutegravir levels were within the normal range; however, the patient’s viral load remained detectable (>50 copies HIV-1 RNA/mL plasma) for the duration of this study (Fig. 1a; 4.5 years).

**Figure 1.**
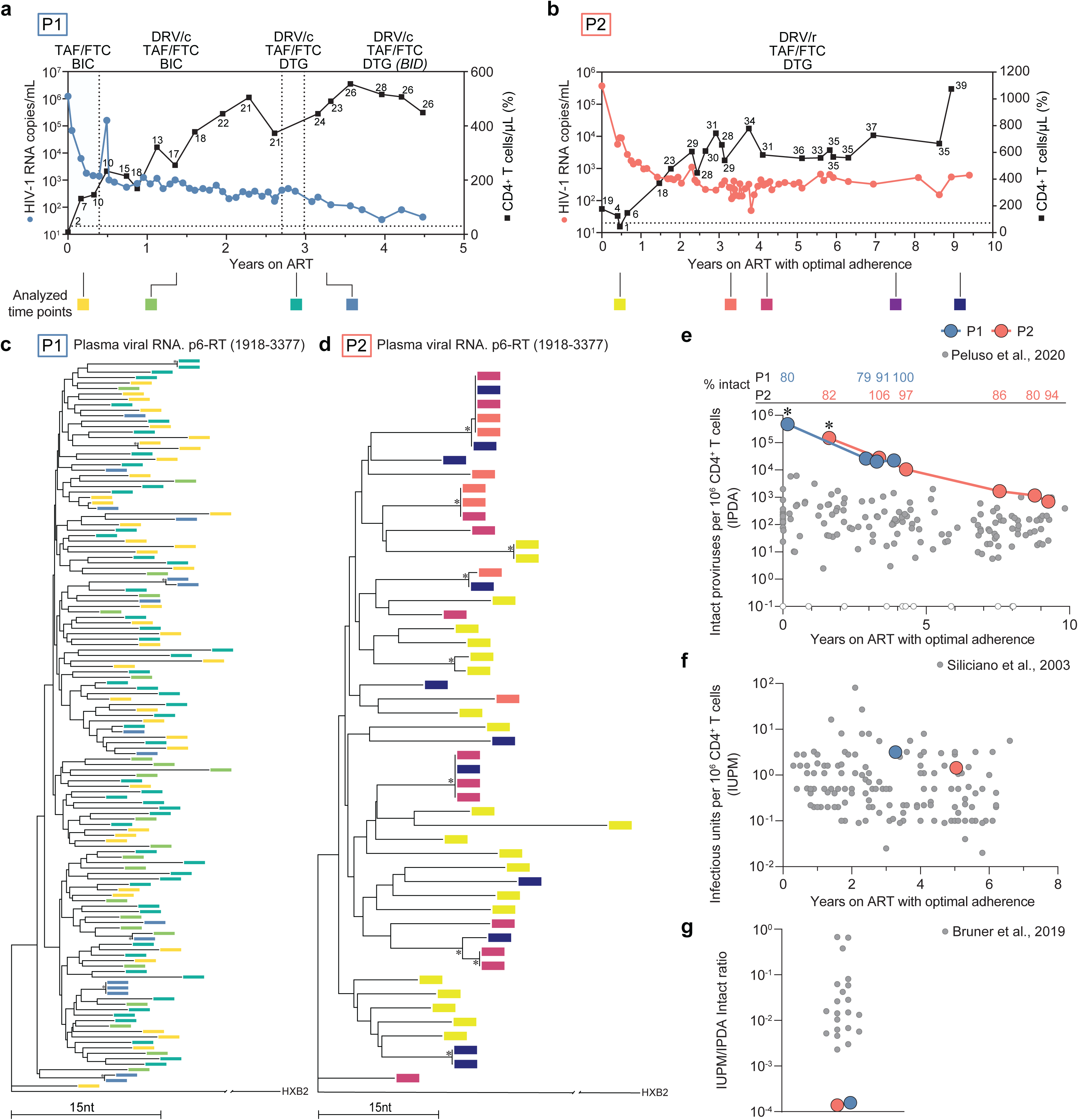
Failure to suppress viremia is characterized by diverse virus in plasma and high infected cell frequency. (**a** and **b**) Plasma HIV-1 RNA and CD4^+^ T cell counts over time for P1 and P2. Numbers above squares represent CD4^+^ T cell percentages. The dotted line at 20 copies/mL represents the current limit of detection for the clinical HIV-1 viral load assay. (**c** and **d**) Neighbor-joining phylogenetic trees of p6-RT single genome sequences obtained from plasma viral RNA for P1 and P2. Phylogenetic tree tip labels are color coded according to the plasma collection timepoint in **a**. Phylogenetic trees are rooted to HXB2 and HIV-1 coordinates refer to the HXB2 reference genome. Tree nodes with bootstrap values above 80 are marked with asterisks. (**e**) Intact proviral DNA frequences as measured by the IPDA. Open circles represent samples where no intact proviruses were detected. The percentage of proviruses classified as intact by the IPDA is shown on the top. Asterisks represent values from PBMC samples which were adjusted based on CD4^+^ T cell percentage. (**f**) Infectious unit per million (IUPM) CD4^+^ T cells as measured by the quantitative viral outgrowth assay. (**g**) IUPM to IPDA intact ratio as measured by dividing the IUPM by the closest IPDA timepoint value. TAF: tenofovir alafenamide; FTC: emtricitabine; BIC: bictegravir; DRV/c: darunavir/cobicistat; BID: *bis in die*; DRV/r: darunavir/ritonavir; DTG: dolutegravir

Participant 2 (P2) is an African American female diagnosed with HIV-1 infection in 2003, with a viral load of 23,343 copies/mL and a CD4^+^ T cell count of 301 cells/μL. P2 initiated ART with atazanavir, lamivudine, and tenofovir disoproxil fumarate one year after diagnosis, resulting in a temporary period of viral suppression (viral load below the limit of detection) (Fig. S1a). Despite multiple attempts at treatment optimization, P2 experienced intermittent adherence to therapy over the next several years, resulting initially in selection for drug resistance, followed by years of non-adherence with concomitant virological failure and an extended period of immunosuppression (Fig. S1a; CD4 nadir of 4 cells/μL). Following hospitalization due to opportunistic *Mycobacterium intracellulare* infection with severe retroperitoneal lymphadenopathy, a new regimen of darunavir, ritonavir, dolutegravir, tenofovir alafenamide, and emtricitabine was initiated (Fig. 1b). Despite consistent adherence to the new regimen for more than nine years, virologic suppression was not achieved during the duration of this study, a surprising finding given that P2 achieved undetectable viremia with the first ART regimen of atazanavir, lamivudine, and tenofovir disoproxil fumarate (Fig. S1a).

Both participants experienced recovery of their CD4 counts (peak CD4 count of 555 and 1,074 cells/µL, respectively) despite the low CD4 nadirs (Fig. 1a, b). Neither participant has an HLA type known to be associated with elite control or rapid disease progression. Since most PWH can achieve viral suppression (<50 copies HIV-1 RNA/mL plasma) within 12 weeks of ART initiation^56–59^, the persistence of detectable viremia after years of treatment with optimal adherence represents a rare and troubling clinical outcome.

### Virus populations in plasma are diverse and show no evidence of evolution

To investigate the cause of this non-suppressible viremia (NSV), we recovered longitudinal single-genome sequences from plasma HIV-1 RNA during ART. Analysis of the HIV-1 *pro-pol* region (1.2 kb amplicon, p6 of *gag* gene through the first 700 bp of reverse transcriptase (RT) in *pol* gene) revealed that the plasma virus sequences were highly heterogeneous (Fig. 1c, d). In P1, 135 out of 139 plasma p6-RT sequences were unique; in P2, 35 out of 51 plasma p6-RT sequences were unique. Sequencing of the *env* gene in P2 confirmed that the plasma virus was polyclonal (Fig. S1b). These results contrast with previous reports of residual viremia following extended periods of suppression which showed that the viremia is typically dominated by few predominant plasma clones^33,36,40–42^. Plasma virus sequences from P1 and P2 lacked the temporal structure characteristic of ongoing viral evolution^23^. Additional analyses showed no accumulation of diversity, divergence, or shift in viral populations over time (Fig. S1c and Methods). Notably, we found no drug-resistant mutations affecting the concurrent ART regimen, a result confirmed through clinical genotyping. These results confirm that NSV was not driven by ongoing replication or selection for drug resistance during ART.

### Large pool of infected cells contributes to persistent viremia

Given the heterogeneity of viral populations in plasma, we hypothesized that consistently detectable viremia in the absence of ongoing replication might reflect the presence of an unusually high frequency of infected CD4^+^ T cells^60^. This hypothesis was supported by the intact proviral DNA assay (IPDA)^61^, which showed that the frequency of proviruses classified as intact in P1 and P2 was markedly higher than previously reported in 81 PWH on ART (151 copies per 10^6^ CD4^+^ T cells; Fig. 1e)^62^. The proportion of proviruses classified as intact by the IPDA at the final time point was 100% and 94% for P1 and P2, respectively – significantly higher than the 6.3% intact proviruses reported in previous studies of PWH on ART (Fig. 1e; P1: *P* = 2.39e-11; P2: *P* = 4.05e-10)^18,63^. We next performed quantitative viral outgrowth assays (QVOAs) to find the frequency of inducible, replication-competent proviruses in P1 and P2. Despite the high frequency of intact proviruses by IPDA (Fig. 1e), the frequencies of cells with inducible, replication-competent proviruses were comparable to previous studies: 3.18 infectious units per million (IUPM) CD4^+^ T cells for P1 and 1.44 IUPM for P2. (Fig. 1f)^11^. The discrepancy between the IPDA and QVOA values is highlighted by the ratio of IUPM to intact proviruses – a metric used to estimate reservoir inducibility^64^ – which was much lower than previously reported in PWH on ART (Fig. 1g)^61^. These results suggest that a much smaller than normal fraction of the proviruses detected in the IPDA could be induced to produce infectious virus.

### The vast majority of infected cells harbor proviruses with shared deletions in key HIV-1 genes

To explain the discrepancy between IPDA and QVOA reservoir measurements, we sequenced near-full length proviral genomes from P1 and P2. Proviruses belonged to subtype B, lacked drug resistance mutations to the concurrent ART regimen, and were CCR5-tropic based on co-receptor prediction algorithms^65^. However, in each participant, greater than 90% of the proviral sequences shared identical large deletions but, surprisingly, were divergent elsewhere in the genome (Fig. 2).

**Figure 2.**
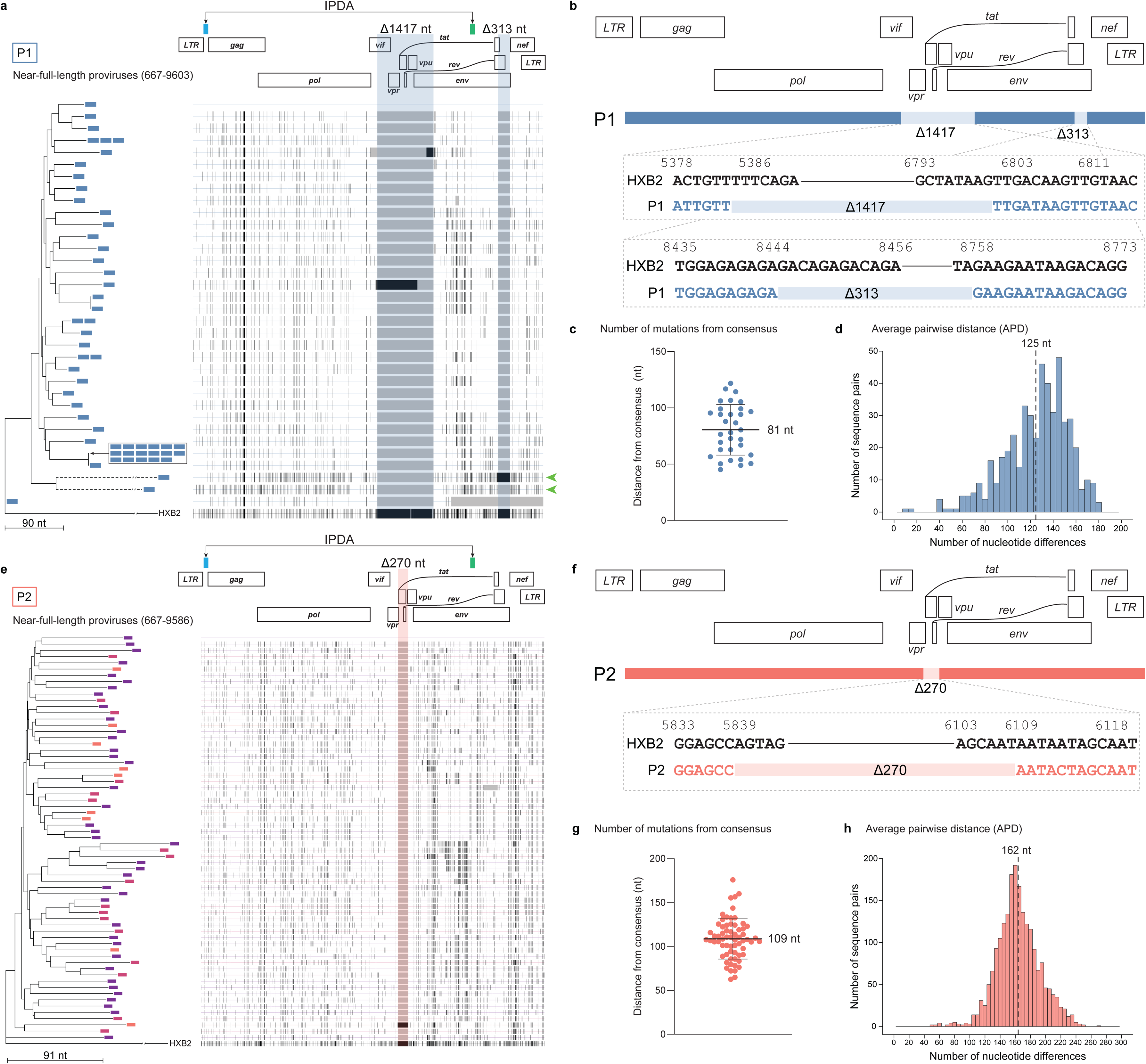
Near-full length sequencing reveals dominant proviral populations with unique deletion signatures and diverse mutations across the viral genome. (**a** and **e**) (Left) Neighbor-joining phylogenetic trees of near-full length proviral sequences obtained by single genome sequencing from primary CD4^+^ T cells, rooted to HXB2. The color of each branch tip indicates sampling time as in Figure 1a and 1b. Dashed branches indicate sequences with hypermutation. (Right) Highlighter plot with black lines representing nucleotide changes compared to the top sequence. Gray vertical bars represent deletions compared to HXB2. Highlighted areas represent recurrent deletion patterns of 1417 nt, 313 nt (P1), and 270 nt (P2). IPDA primer probe regions are highlighted at the top. Green arrows point to sequences with significant G -> A hypermutation. (**b** and **f**) Mapped sequences of prominent deletion signatures found in majority of proviruses in P1 and P2 compared to HXB2. (**c** and **g**) Dot plots representing the number of mutations between each near-full length sequence and the majority consensus sequence. Horizontal line represents mean, error bars represent SD. (**d** and **h**) Histogram (bin width of 5) representing the nucleotide differences in unique proviruses from each participant. Dashed line represents the mean number of nucleotide differences between all the unique proviral sequences.

In P1, 50 out of 54 sequences contained the same set of two deletions of 1417 and 313 nucleotides in exactly the same positions. The 1417 nt and 313 nt deletions affected the open reading frames of *vif, vpr, tat, rev, vpu,* and *env* (Fig. 2a, b). The 1417 nt deletion eliminated *vpr* and *vpu* genes, the first exons of *tat* and *rev*, and the first 578 nucleotides of the *env* gene, likely preventing expression of these proteins. Proviruses with these deletions were classified as defective according to the NCI ProSeq-It tool^66^. The nature of these deletions would almost certainly preclude further replication, yet the proviruses harboring these exact two deletions were remarkably diverse elsewhere in the genome (Fig. 2a). To assess the diversity within the proviral sequences, we employed two different approaches. We first compared the number of nucleotide changes between each provirus to a majority consensus sequence to estimate the extent of viral diversification. A consensus sequence of all proviruses with the deletions provides a way to infer the sequence of the provirus in the cell in which the deletions initially arose. The mean distance of the proviral sequences to the majority consensus sequence was 81 nucleotides (Fig. 2c). Next, we measured the average pairwise distance to find the genetic variation within the proviral population. The average pairwise distance of the unique near-full length proviral sequences was 1.38% suggesting that, on average, there were 125 nucleotide differences between each unique sequence (Fig. 2d). Furthermore, some of the proviruses with the recurring deletions also had signatures of APOBEC3G/F-mediated hypermutation (Fig. 2a, dashed phylogenetic tree branches and green arrows), indicating that viral RNA harboring these deletions entered a target cell where it underwent hypermutation during reverse transcription. Only one proviral sequence matched the plasma virus in the p6-RT regions, reflecting the extreme diversity and size of the infected cells population.

In P2, we found a 270-nucleotide deletion impacting the reading frames of *vpr, tat, rev,* and *vpu* in 64 out of 65 sequences (Fig. 2e, f). Given the critical roles of Tat and Rev in the virus life cycle, it is expected that this deletion would preclude replication of the deleted virus. Nevertheless, as with P1, proviruses with this deletion showed significant diversity elsewhere in the viral genome. The mean distance from the majority consensus sequence was 109 nucleotides (Fig. 2g). The average pairwise distance of the near full length proviral sequences with this deletion was 1.81% (Fig. 2h), corresponding to an average of 164 nucleotide differences among proviruses. The deletions found in proviruses from P1 and P2 did not impact the binding of the IPDA primers and probes (Fig. 2a, e), explaining the high frequency of proviruses classified as intact (Fig. 1e).

Identical sequences harboring fatal deletions typically indicate proliferation of CD4^+^ T cell clones carrying defective proviruses^31,33,40,67–69^, since these deletions render the virus unable to replicate^17,18,36^. However, analysis of the near-full length proviral sequences in P1 and P2 suggests that the high diversity between each defective provirus cannot be attributed to clonal expansion, errors made by human RNA polymerase II, or PCR and sequencing errors. Instead, the finding of nucleotide differences in the viral genome that carry exactly the same large fatal deletions implies that the defective viral RNA was packaged into virions, entered a target cell, underwent reverse transcription leading to additional HIV-1 RT-mediated mutations, and was then integrated into the host genome.

To specifically quantify proviruses harboring these deletions, we designed custom digital PCR assays with probes spanning the deletion junction and primers bracketing the deleted region (Fig. 3a). For P1, we designed a triplex assay that quantifies each deletion separately in addition to the region upstream *gag* (LTR-gag). For P2, we designed a competition assay that distinguishes proviruses with or without (WT-270) the 270 nt deletion (Fig. 3a, b). Assay specificity was validated by testing each assay on genomic DNA isolated from other PWH; only DNA from P1 and P2 yielded positive signals from the deletion-specific PCRs (Fig. S2). We found that greater than 80% of proviruses contained these deletions, and the proportion of proviruses containing the deletions remained stable over time on ART (Fig. 3c). Taken together, these results show that most infected cells contained these deletions, suggesting a unique situation that allowed their accumulation during untreated infection and their persistence over time on ART.

**Figure 3.**
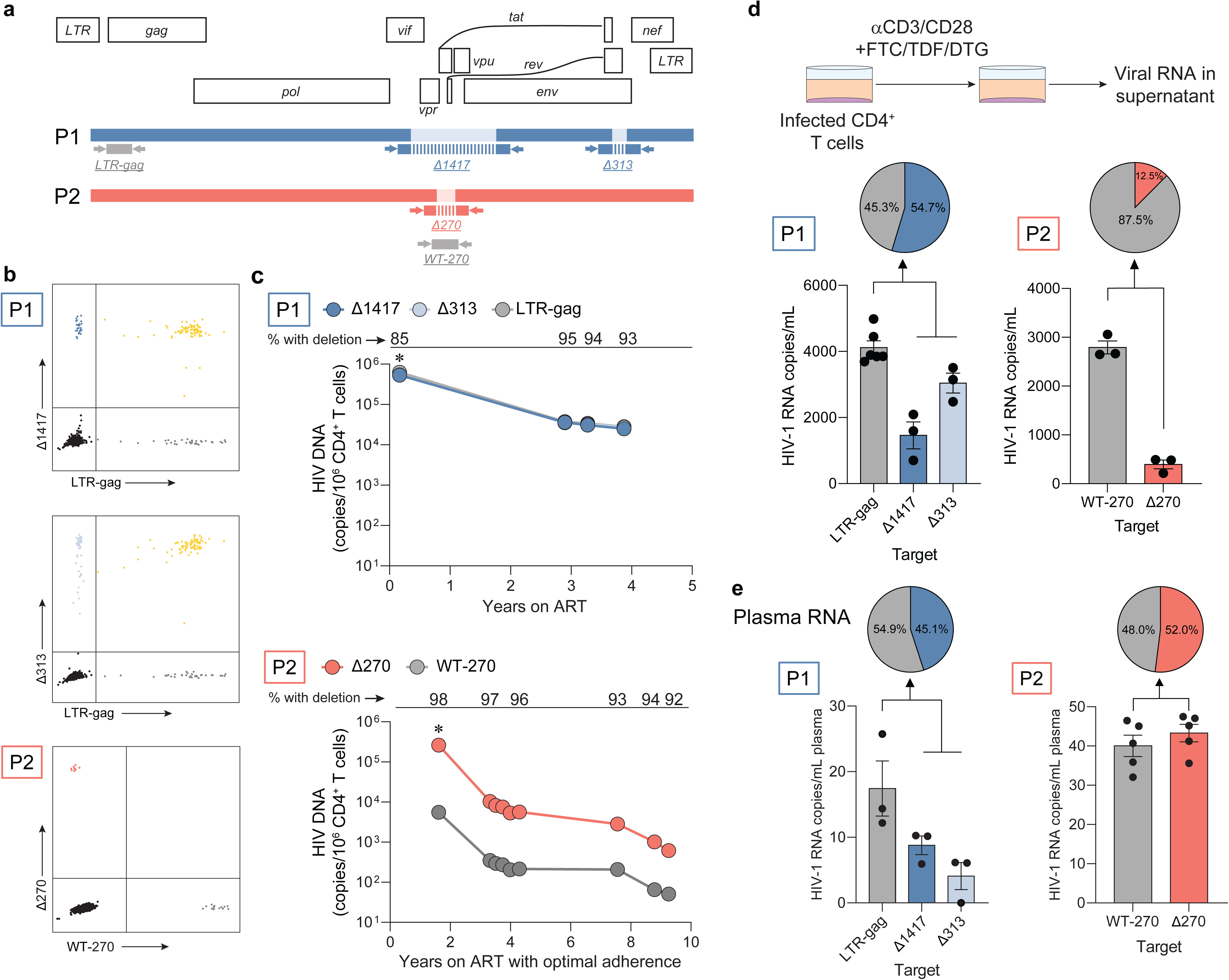
Defective proviruses dominate the proviral landscape and contribute to NSV. (**a**) Location of primers (arrows) and probes (rectangles with vertical bars) to specifically quantify deletion signatures in viral RNA and DNA. Probes span the deletion region. (**b**) Representative two-dimension plots of dPCR showing duplex amplification of intact proviruses and proviruses of interest by deletion-specific assays. (**c**) Longitudinal quantification of proviruses with specific deletions of interest. (**d**) CD4^+^ T cells from P1 and P2 were cultured for 48 or 72 hours in the presence of emtricitabine (FTC), tenofovir disoproxil fumarate (TDF), dolutegravir (DTG) and anti-CD3/CD28 beads. The virion-associated RNA in the supernatant was measured by RT-dPCR. Error bars represent the SEM. Pie chart shows percentage of HIV-1 RNA copies with deletion normalized to the total number of copies. (**e**) Plasma virion-associated RNA was measured by RT-dPCR. Error bars represent SEM. Pie chart shows percentage of HIV-1 RNA copies with deletion normalized to the total number of copies. For P1, total proviruses were quantified by measuring the highly conserved region LTR-gag. For P2, proviruses without the 270 nt deletion were measured by using a primer-probe set inside of the deletion (WT-270).

### Proviruses with recurrent deletions can express viral RNA and contribute to persistent viremia

Given that some defective proviruses can express HIV-1 RNA and produce viral proteins *in vivo*^36,38,39^, we evaluated whether proviruses harboring these fatal deletions could be induced to produce virions that package the defective viral RNA. We stimulated CD4^+^ T cells from P1 and P2 with anti-CD3/CD28 beads in the presence of antiretroviral drugs (TDF, FTC, DTG) and quantified viral RNA in the supernatant using a digital RT-PCR assay with the same primers and probes described above. In P1, we found that, on average, 55% of virion-associated RNA contained either the 1417 nt or 313 nt deletion (Fig. 3d). Due to shearing during the RNA isolation, it was not possible to quantify the viral RNA that contained both deletions. Differences between the frequencies of the 1417 and 313 nt deletions can be explained by mutations surrounding the deletions which prevent primer or probe binding or viral genomes which only had one of the deletions. In P2, 13% of the virion-associated RNA in the supernatant contained the 270 nt deletion (Fig. 3d).

Given that infected cells from both participants could be induced to make virions packaging these defective RNAs, we determined if cells with these proviruses contributed to the NSV. Using the same digital PCR assays, we found that in both participants, about 50% of virion-associated HIV-1 RNA in plasma harbored these deletions (Fig. 3e).

We also analyzed QVOA isolates to determine whether proviruses with these specific deletions could be induced to produce infectious virus. Despite an abundance of cells carrying defective proviruses seeded into each outgrowth well, the majority of isolates that exponentially grew out in the culture were composed of virions packaging intact viral genomes (Fig. S3). However, some wells with outgrowth also contained virions packaging defective viral genomes in addition to virions with intact genomes, consistent with our results from *ex vivo* stimulation of infected CD4^+^ cells and analysis of plasma viremia.

Together, these results show that although the recurring deletions are incompatible with replication, the viral RNA from the dominant deleted proviruses could still be packaged into virus particles and contribute to persistent viremia.

### Molecular clones with these deletions show significantly reduced virion production

To further investigate the impact of deletions on replicative fitness and virus production, we introduced the recurring deletions into a reference proviral construct (NL4-3) (Fig. 4a)^70^. Next, we transfected HEK293T cells and measured virion production after 72 hours. As expected, deletions in key HIV-1 genes resulted in an almost complete loss of virus production (>100,000-fold reduction) compared to the wild type, as measured by p24 ELISA (Fig. 4b). We then stained the surface of the HEK293T cells with multiple broadly neutralizing antibodies (bNAbs) to assess Envelope (Env) expression (Fig. 4C). Given that the 1417 nt deletion impacts V1 and V2 of *env*, we tested bNAbs targeting V1/V2-dependent and independent epitopes (Fig. S4). Consistent with the p24 ELISA results, transfection with the defective molecular clone resulted in a dramatic reduction in the percentage of HEK293T cells expressing Env, to levels that were not statistically different from cells transfected with a construct lacking a functional *env* gene (NL4-3-ΔEnv, n.s. *p* > 0.01; Fig. 4d).

**Figure 4.**
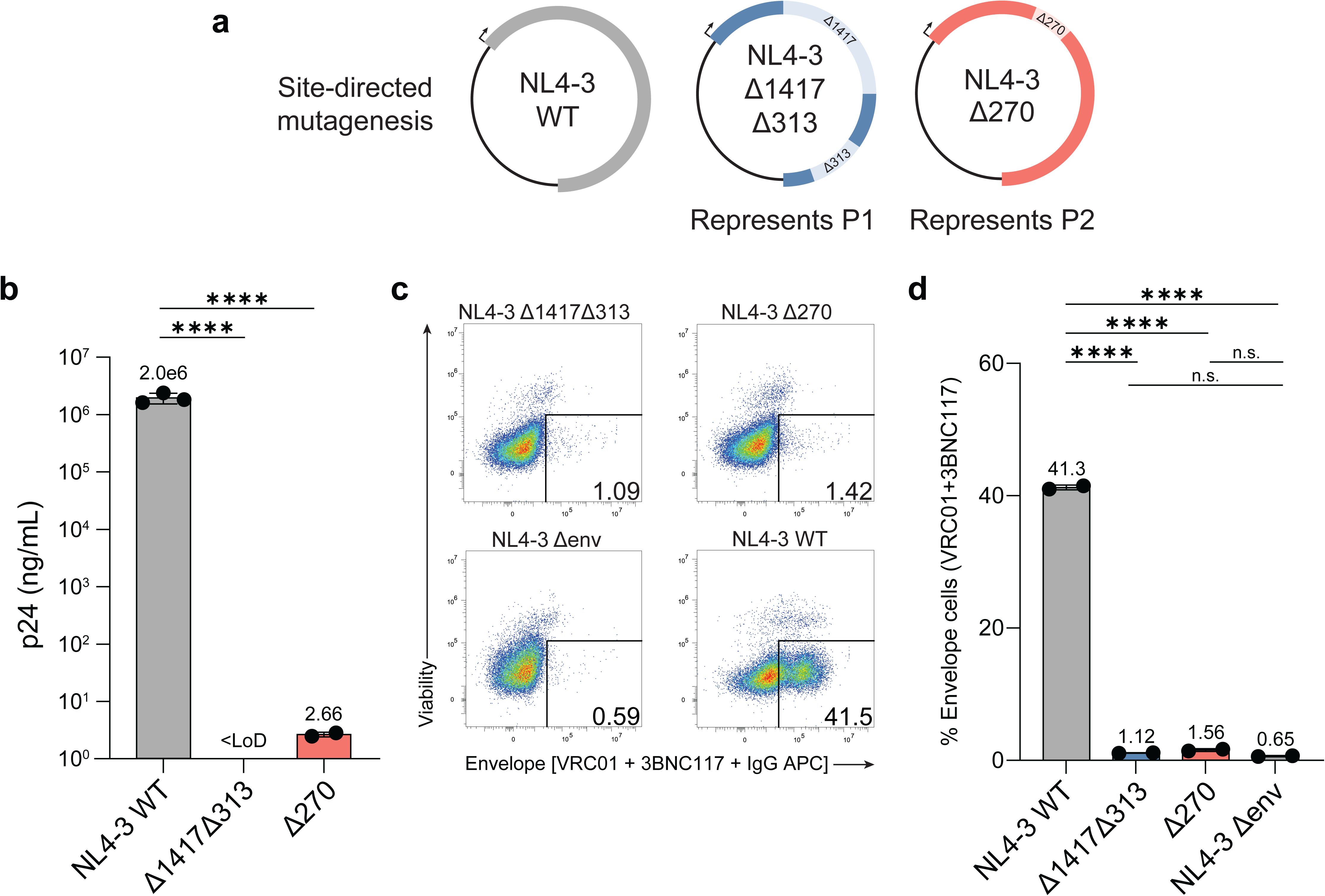
Deletions found in proviruses abolish virus production *in vitro*. (**a**) Prominent deletions found in proviruses were introduced into an NL4-3 expression plasmid by site-directed mutagenesis. Arrows represent orientation of the provirus within the expression plasmid. (**b**) Virus produced upon HEK293T transfection was pelleted by ultracentrifugation, and p24 was measured by ELISA. Lower limit of detection (LoD) was 0.625 ng/mL. Horizontal bars represent mean, and error bars represent SD (**c**) Representative flow cytometry plots of the surface staining of transfected HEK293T cells with viability dye and bNAbs (VRC01 and 3BNC117). (**d**) Surface staining of HIV-1 Env with bNAbs (VRC01 and 3BNC117) on HEK293T cells 24 hours after transfection. Horizontal bars represent mean and error bars represent SEM. (**b** and **d**) Statistical significance between conditions was determined by 1-way ANOVA. **** *P* < 0.0001. n.s. *P* > 0.01

These observations show that virion-associated RNA in the supernatant of activated infected cells and in plasma contained the same deletions found in the proviral sequencing. However, the deletions found *in vivo* resulted in abrogated virion production when introduced into replication-competent molecular clones. To address this apparent discrepancy, we investigated alternative mechanisms for the dissemination of these proviruses.

### Dissemination of defective proviruses via superinfection

We hypothesized that intact HIV-1 virions could superinfect cells carrying proviruses with the recurrent deletions and drive the production of infectious viral particles carrying defective genomic RNA. In superinfected cells, viral proteins can be produced from the intact provirus, but genomic RNA from both defective and intact proviruses compete for packaging into the virions^71^. This allows infectious particles that have packaged the defective viral RNA to infect new cells, where the defective RNA genome undergoes reverse transcription, accumulates additional mutations, and then integrates into the host cell genome. In principle, the host cell could then be superinfected by another intact virus, leading again to mobilization of the defective genome and accumulation of additional mutations in the next round of infection. In this manner, defective proviruses carrying the same fatal deletions could accumulate a diverse set of additional mutations. The competition between replication-competent and defective viral genomes for packaging in co-infected cells could result in viral interference, consistent with DIPs^44,50^. Therefore, we hypothesized that viruses with the recurrent deletions could interfere with WT (intact) virus replication. To assess this hypothesis, we generated NL4-3-based reporter construct (NL4-3-ΔNef-BFP) which harbored the deletions found in each participant (P1: Δ1417Δ313; P2: Δ270). We created model cell lines for each participant by transducing SupT1 cells at low multiplicity of infection and bulk sorting BFP^+^ cells for downstream assays (Fig. S5a). To assess the ability of the deleted-variants to interfere with replication of WT virus, we infected the model cell lines with an infectious WT reporter NL4-3 virus carrying RFP in the *nef* ORF (NL4-3-ΔNef-RFP) and measured the frequency of cells infected with the WT virus (RFP^+^) over time (Fig. 5a, b, Fig. S5b). The spread of WT virus in cells carrying the defective proviruses was significantly reduced compared to WT virus spreading in mock-transduced SupT1 cells, suggesting that these defective proviruses interfere with WT virus replication (Fig. 5b). We quantified the infectivity of WT virus by using the supernatant from day 3 of culture to infect new SupT1 cells in a single-round infectivity assay (Fig. 5c)^72^. WT virus infectivity was significantly reduced when using the supernatant from the deleted-variant cell lines compared with the mock-transduced cell line, further supporting the conclusion that the defective proviruses interfere with WT virus replication (Fig. 5c). WT viral titers were similarly reduced in both model cell lines carrying the defects observed in P1 and P2 (Fig. 5c). To test whether the superinfected cells produced infectious virions packaging either the intact or defective RNA, we quantified the percentage of RFP^+^ (WT virus) or BFP^+^ (defective virus) SupT1 cells after a single round of infection. When infected with supernatant from the superinfected cells, >40% of target cells were infected with the defective virus (BFP^+^) (Δ1417Δ313: 43%; Δ270: 62%) (Fig. 5d).

**Figure 5.**
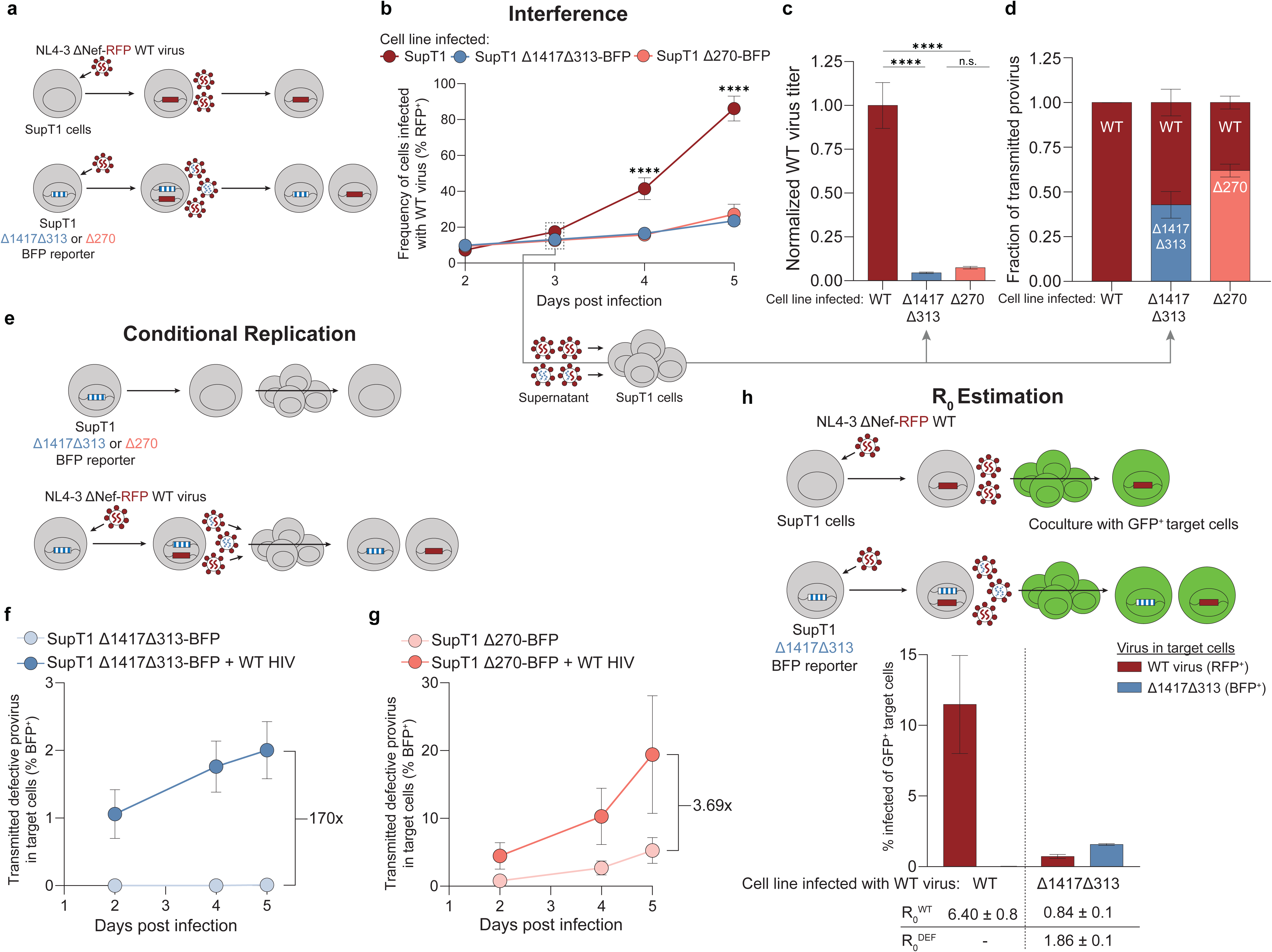
Cell culture model to assess superinfection *in vitro*. (**a**) Schematic for assessment of superinfection *in vitro*. (Top) SupT1 cells are infected with NL4-3-ΔNef-RFP WT virus. (Bottom) Model cell lines transduced with reporter defective viruses representing P1 (Δ1417Δ313) or P2 (Δ270) are infected with NL4-3-ΔNef-RFP WT virus. Virions produced by these superinfected cells may package RNA from the defective (blue) or WT (red) viral genome. (**b**) Frequency of cells infected with the WT virus (% RFP^+^) measured over time. Points represent mean and error bars represent SD. (**c**) Viral supernatant from day 3 of infection of each cell line was used to infect target SupT1 cells to determine the viral titer of WT virus after single round of infection. WT viral titer was normalized to virus from the WT SupT1 cell line. Horizontal bars represent mean and error bars represent SEM. (**d**) Viral supernatant from day 3 of infection of each cell line was used to infect SupT1 cells to determine the fraction of transmitted provirus after single round of infection as measured by flow cytometry. (**e**) Schema for assessment of conditional replication *in vitro*. Model cell harboring reporter defective viruses are either mock-infected or infected with WT virus. After 3 days of culture, the resulting supernatant is used to infected target SupT1 cells. (**f** and **g**) Frequency of target SupT1 cells infected with defective provirus measured over time. Points represent mean and error bars represent SD. (**h**) (Top) Schema for three-color experiment to estimate R_0_ *in vitro*. WT SupT1 or SupT1-Δ1417Δ313-BFP are infected with WT virus. After 2 days, the cells were cocultured with excess GFP^+^ SupT1 target cells. (Bottom) Frequency of target GFP^+^ SupT1 target cells infected with either WT virus (red) or defective virus (blue). R_0_ values and standard error of mean for both the WT virus (R ^WT^) and defective virus (R ^DEF^) are listed below. Horizontal bars represent mean and error bars represent SD. (**b** - **h**) n=6 for all conditions assessed. (**b** - **c**) Statistical significance between cell lines was determined by 1-way ANOVA. **** *P* <0.0001. n.s. *P* > 0.05.

Next, to validate that the defective proviruses conditionally replicate in the presence of a WT virus, we collected supernatants from either the deleted-variant cell lines alone or from the deleted-variant cell lines which had been infected with WT virus for 3 days. These supernatants were used to infect SupT1 cells. We then assessed the target SupT1 cells for the presence of defective proviruses (BFP^+^) (Fig. 5e). For the cell line representing P1, the frequency of defective proviruses in the target cells increased by >170-fold when using viral inoculum from superinfected cells, compared to inoculum from cells which were not superinfected with WT virus. For the cell line representing P2, the frequency of defective proviruses increased 3.69-fold when target cells were infected with viral inoculum from cells that were superinfected, compared to inoculum from cells which were not superinfected (Fig. 5e). These data show that these defective proviruses transmit their genome to target cells when superinfected with WT virus. Together, these results demonstrate that the recurrent proviral deletions found in P1 and P2 interfere with WT virus replication and conditionally replicate only in the presence of WT virus, two hallmarks of DIPs^73^.

Given that the deletions found in P1 (Δ1417Δ313) strongly resembled the engineered deletions which constitute the therapeutic interfering particle (TIP) described in Pitchai, Tanner et al. (e.g., deletion of *tat*, *rev*, *vpu*, *env*, and retention of cPPT) (Fig. S5c), and considering the ability for viral genomes harboring this deletion to integrate into target cells (Fig. 5d), we hypothesized that the defective proviruses found in P1 functioned as a TIP^50^. For a DIP to be considered a TIP, it must be mobilized with an R_0_ > 1, meaning that the defective provirus can give rise to virions that infect more than one target cell and therefore spread *in vivo*. To directly measure the R_0_ of the defective variants (R_0_^DEF^), we implemented a three-color assay (Fig. 5h)^50^. First, we infected the Δ1417Δ313-BFP reporter cell line with the WT-RFP virus as in Figure 5b. After 2 days, we cocultured 30,000 infected cells with 70,000 GFP^+^ SupT1 target cells (see Methods for generation of the GFP^+^ target cell line). After 3 days of co-culture, we assessed the fraction of GFP^+^ target cells with either the WT virus (RFP^+^) or the defective virus (BFP^+^), allowing us to estimate the frequency of secondary infection events (Fig. S5d). The R_0_ was found by dividing the number of secondary infections (i.e., infected target cells) by the number of primary infections (i.e., initial infection events) (see Methods). The R_0_^WT^ following infection of wildtype SupT1 cells with WT virus alone was 6.40, consistent with lower estimates in previous reports^74–76^. As expected, the R_0_^WT^ in the Δ1417Δ313 reporter cell line (R_0_^WT^: 0.84) was significantly less than the R_0_^WT^ in WT SupT1 cells, consistent with viral interference (Fig. 5b; *p* = 0.0002). The R_0_^DEF^ represents the average number of target cells infected with the defective virus resulting from WT virus superinfection of cells carrying the deleted provirus. Notably, the R_0_^DEF^ in the Δ1417Δ313 reporter cell line was 1.86 (Fig. 5h). This result is consistent with the spread of the defective viruses *in vivo*. The low R_0_ of the defective lineage suggests that although viral interference can be detected *in vitro* (Fig. 5b), the interference does not affect WT viral replication and pathogenesis *in vivo* to a clinically significant extent, as evidenced by P1’s high initial viral load and extremely low CD4 nadir at the time of ART initiation.

### Superinfected CD4^+^ T cells carrying both intact and defective proviruses can be detected ex vivo

To show that defective proviruses found in P1 and P2 were mobilized – before the introduction of effective ART – by superinfection with intact proviruses, we looked for cells carrying multiple proviruses in primary CD4^+^ T cells from P1 and P2. We modified the HIV-Flow assay^77^ to single-cell sort p24^+^ cells into individual wells and lysed the cells (Fig. 6a). Similar to previous work investigating the number of proviruses within individual infected cells^78^, we next distributed the cell lysate over 6 or 12 wells, resulting in a greater than 80% probability that, if two proviruses were present, they would be separated into different wells (Fig. 6a, Fig. S6a). Lastly, we amplified and sequenced the near-full length proviral genomes^79^. Given the importance of sorting accuracy in this experiment, we first sorted individual CD4^+^ T cells from an HIV-1 negative male donor and confirmed that all sorted cells were singlets by amplification of the *SRY* gene, which is found in a single copy per cell on the Y chromosome (Fig. S6b; methods). We also validated this experimental protocol by sorting p24^+^ ACH-2 cells. In accordance with other reports, we found that ACH-2’s can harbor multiple proviruses per cell when cultured in the absence of antiretroviral drugs (Fig. S6c)^80–82^. We further validated this protocol using cells from a participant with low-level viremia: out of 5 sorted p24^+^ cells, none had evidence of containing more than one provirus, consistent with previous findings^78,83,84^ (Fig. S6d; methods). In a recent study, Dufour and colleagues did not find any evidence of cells infected with multiple proviruses from 305 single-sorted p24^+^ cells from 6 PWH on suppressive ART^79^ (Nicolas Chomont, Caroline Dufour, personal communication).

**Figure 6.**
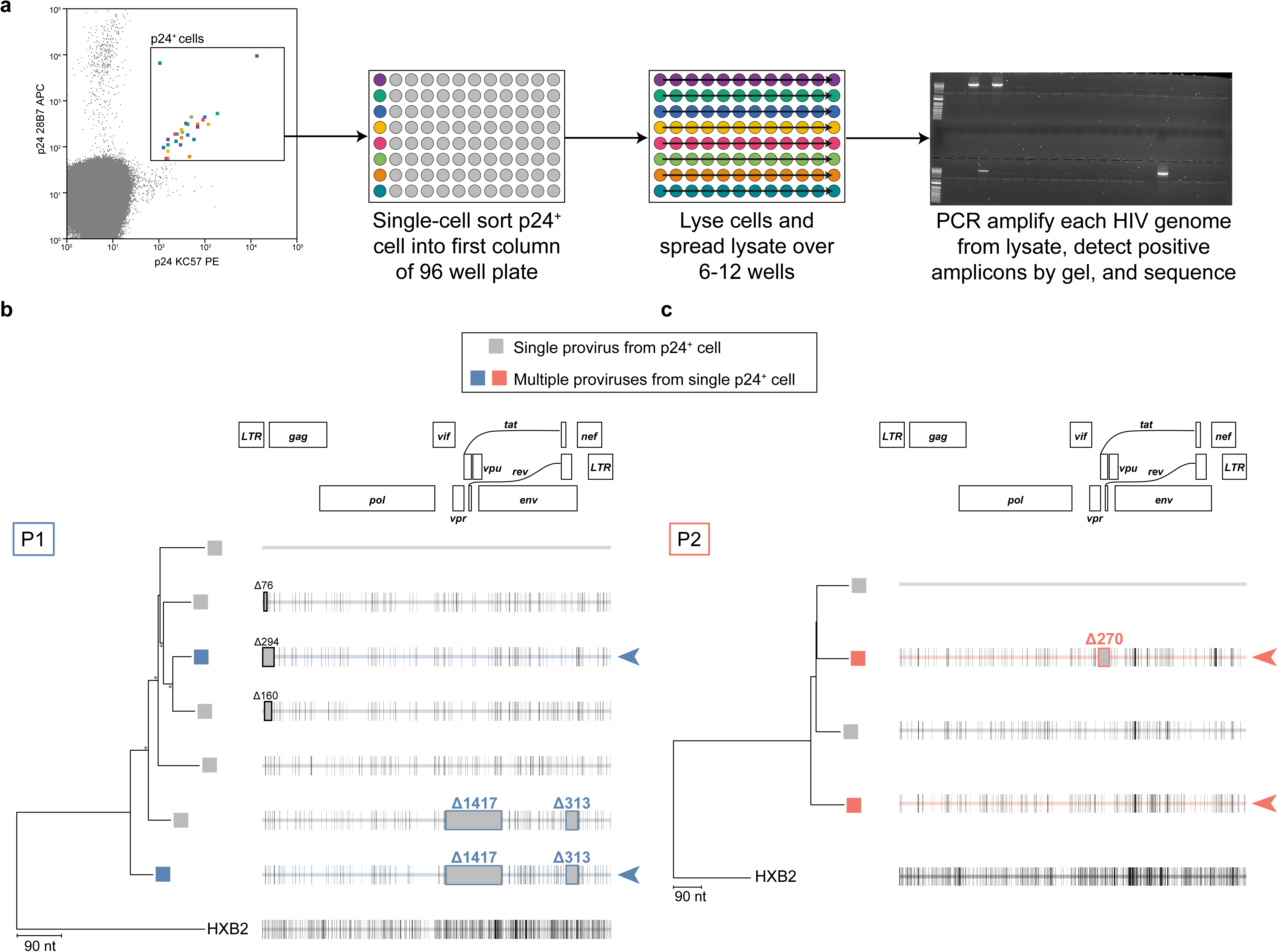
Single p24^+^ cell sequencing identifies dually infected cells *in vivo*. (**a**) Experimental design to characterize p24^+^ cells and detect multiple integrated proviruses. Sorted p24^+^ cells are lysed and spread over multiple wells. Near-full length proviral amplification, agarose gel electrophoresis, and sequencing are conducted on the cell lysate. Example agarose gel electrophoresis shown here. (**b** and **c**) Neighbor-joining phylogenetic tree of near-full length proviral genomes from p24^+^ cells. Highlighter plot with black lines representing nucleotide changes compared to the top sequence. Gray boxes represent deletions compared to the top sequence of each tree. Tree nodes with bootstrap values above 80 are marked by asterisks. 5’ internal deletions are boxed in black and annotated. Common deletions found in P1 and P2 are boxed and annotated. Phylogenetic tree tips with gray boxes represent single provirus found from a sorted cell. Colored boxes represent multiple proviruses found from a single-sorted cell, likely representing multiple integrated proviruses within one cell. Colored arrows point to multiple proviruses found in a single-sorted cell.

For P1, we analyzed ∼13 million CD4^+^ T cells and recovered 7 sequences from the p24^+^ sorted cells (0.54 proviruses per 10^6^ CD4^+^ T cells) (Fig. 6b). From two of the p24^+^ sorted cells, we identified proviruses that had small deletions in the 5’ region of the viral genome – which are disproportionally found in p24^+^ cells^79,85,86^ – yet did not have the 1417 nt and 313 nt deletions found in the bulk proviral population (Fig. 2a). From another two of the p24^+^ sorted cells, we identified intact proviruses. Notably, we identified one cell which had two integrated proviruses: one provirus with the 1417 nt and 313 nt deletions and one nearly intact provirus with a small 5’ deletion, demonstrating the presence of superinfected cells *in vivo* (Fig. 6b, color boxed). Lastly, from another sorted cell, we identified one provirus harboring both the 1417 nt and 313 nt deletions, a surprising finding given that these two deletions decrease p24 production by several orders of magnitude *in vitro* (Fig. 4b). Given the low efficiency of near-full length proviral amplification^87^, it is possible that we detected only one of the two proviruses in that cell. For P2, we similarly analyzed ∼13 million CD4^+^ T cells and recovered 4 sequences from the sorted cells (0.30 proviruses per 10^6^ CD4^+^ T cells) (Fig. 6c). From two cells, we identified one intact provirus in each. Notably, in one p24^+^ sorted cell, we detected two proviruses – one carrying the 270 nt deletion and one that was intact across the entire viral genome (Fig. 6c, colored boxes).

Together, these results provide direct evidence of persistent infected cells carrying two proviruses and support our *in vitro* studies showing that superinfection allowed the mobilization of defective proviruses and the accumulation of mutations.

## Discussion

Most PWH achieve suppression of plasma HIV-1 RNA to below the limit of detection (50 copies HIV-1 RNA/mL plasma) within 12 weeks of ART initiation^56–58^. However, HIV-1 persists in a latent reservoir in resting CD4^+^ T cells^5,6^, and daily reactivation of a small fraction of these infected cells gives rise to residual viremia^40,88–90^. In rare cases, PWH experience detectable viremia after years of suppression despite adherence to ART^33,36,41,42^. In this situation, the viremia cannot be reduced by treatment intensification and is termed non-suppressible viremia (NSV)^91^. NSV is typically due to large clones of infected cells. In contrast, we describe a novel presentation of NSV in which treated PWH never achieved an undetectable viral load despite optimal adherence. Surprisingly, the NSV in these cases reflected a large and diverse reservoir of infected cells that was generated by mobilization of defective proviruses. Here, we provide direct evidence that the defective proviruses found in the study participants function as DIPs, a finding that has important implications for the use of TIPs as a novel therapeutic strategy.

We first interrogated the source of the viremia in participants who never achieved undetectable viral loads and found surprising sequence heterogeneity, suggesting that the NSV was caused by a novel mechanism distinct from previous reports^33,36,41,42^. The proviral frequency in both participants suggested that the reservoir was remarkably large compared to most PWH on ART, with almost every provirus being classified as intact by the IPDA. Near-full length sequencing revealed a very surprising result: almost all proviruses harbored exactly the same large deletions yet had numerous distinct mutations elsewhere in the genome. These deletions did not impact the binding of the IPDA primers and probes, explaining the high frequency of proviruses classified as intact by the IPDA. These defective proviruses could be induced to generate virions packaging defective RNA both *in vivo* and *ex vivo*. However, introducing these deletions into infectious molecular clones abolished virion production and prevented new infection events, suggesting that these deletions rendered the provirus fatally defective. Therefore, we hypothesized that before ART was initiated, cells harboring defective proviruses were superinfected with replication-competent viruses, leading to the incorporation of defective genomes into infectious virus particles that could be released from superinfected cells, resulting in the propagation of the fatal deletions and introduction of new mutations through RT-generated errors in the newly infected cells. Subsequent infection of these cells by replication-competent virus could lead to further rounds of mobilization of the defective genome and accumulation of additional mutations. Overtime, this process could generate a diverse set of defective proviruses, all harboring the same fatal deletions. Given the HIV-1 RT error rate and the single round infectivity of the defective lineages, this process must have occurred for extended time intervals in order to result in the striking diversity we observed. We developed novel cell lines carrying the proviruses with the recurrent deletions and found that the defective proviruses could only propagate when superinfected with wildtype virus. Additionally, the recurrent deletions found in P1 replicated with an R_0_ > 1 when introduced into the reference plasmid, NL4-3, suggesting that viruses with these deletions can propagate when superinfected. Lastly, we found direct evidence of superinfected cells from both participants, suggesting that superinfection was the mechanism responsible for the propagation of defective proviruses.

Although HIV-1 has evolved mechanisms to prevent superinfection (downregulation of CD4 by Nef, Env, and Vpr), superinfection has been documented in spleen tissue from PWH during untreated chronic infection^92–94^. Additionally, superinfection has been implicated in the loss of viral control in some post-treatment controllers^95^. Importantly, superinfection is the major mechanism for viral recombination, contributing to the formation of new circulating strains^96^ and intra-host evolution^97^, including the selection for drug resistance^98^. Although superinfection of cells *in vitro* has been shown to occur with greater than expected frequency^99–101^, multiple studies have shown that >85% of CD4^+^ T cells from the peripheral blood and lymph node tissues in PWH contain only a single viral genome^78,84^. Superinfection is also rare in other human retroviruses. During chronic HTLV-1 infection, the majority of infected cells contain a single provirus^102^. To our knowledge, previous to this work, there is no report of cells harboring multiple proviruses in PWH on ART.

Superinfection has also been investigated in the context of DIPs^48,73,103^. Although DIPs have been shown to arise for most RNA viruses, they have not been found in PWH. This study provides the first evidence for HIV-1 DIPs *in vivo*. HIV-1 DIPs are likely extremely rare; the formation of an HIV-1 DIP requires a very specific type of error during reverse transcription: the defective provirus must retain the packaging/encapsidation signal (Ψ) and the Rev response element (RRE). These two regions are required for defective RNA to be exported from the nucleus and to be packaged into viral particles. As a result, the generation of proviruses which can serve as the “chassis” for DIPs is a rare event that may occur only after years of untreated infection. Additionally, the accumulation of a large number of distinct mutations on the deleted provirus “chassis” likely required a prolonged period of untreated infection, as is documented in the extended clinical history of P2 (Fig. S1a). Given that a pre-therapy clinical history was unavailable for P1, we estimated that 407 cycles of mobilization occurred, requiring a minimum of 2.23 years, based on the error rate of the HIV-1 RT (3.4 x 10^-5^ mutations/bp/cycle) and a 2 day generation time^21,104,105^. This estimate does not take into account the possibility of recombination. Nevertheless, proviruses with the recurrent deletions were likely disseminated and diversified during a prolonged period of high-level viral replication prior to ART. Importantly, the defective proviruses identified in this study likely propagated at a time with low CD4^+^ T cell count (estimated range: 10^1^ - 10^2^ cells/μL), increasing the likelihood of superinfection.

Interestingly, in both participants, the DIPs did not prevent CD4^+^ T cell depletion. Additionally, the putative DIPs did not result in a reduction in setpoint HIV-1 RNA levels, although the clinical viral load measurements cannot discriminate between intact and defective virus. Recent reports have suggested that molecular engineering of DIPs to have an R_0_^DEF^ > 1 can allow them to function as a therapeutic interfering particle (TIP), reducing viremia and pathogenesis^50^. The defective proviruses from P1 appeared to have an R_0_^DEF^ of 1.86 in *in vitro* assays, which was markedly lower than engineered TIPs (∼12) despite their sequence similarity^50^ (Fig. S5c). Despite promising results in pre-clinical studies, the ability of TIPs to reduce viral set point and pathogenesis in PWH is unknown. The very low CD4 nadirs and high viral loads described here raise concerns about the effectiveness of the TIP strategy, but further investigation is clearly required. Our work further shows that, in addition to 5’ leader defects, other types of defective proviruses can contribute to viremia^36,39,42^. These proviruses, if transcriptionally active, may complicate the interpretation of viral load, leading to potential misinterpretations in clinical and cure-focused studies.

Our study has limitations. Principally, we were restricted to peripheral blood samples taken after years on ART and not during the period of untreated infection in which the defective genomes propagated. Future work should aim to further interrogate the prevalence of superinfected cells in PWH with high frequencies of infected cells who are unable to reach an undetectable viral load.

In conclusion, we demonstrate that the failure to achieve an undetectable viral load despite effective ART can be caused by a large, heterogeneous reservoir of infected cells generated by a novel process involving the dissemination of defective proviruses through DIPs. In this situation, the proviral landscape can be dominated by diverse proviruses that share the same lethal deletions. Such a paradox led us to identify superinfection as a novel mechanism for the mobilization of defective proviruses. Although we describe DIPs in PWH for the first time, the DIPs found in both participants did not prevent CD4 T cell depletion, raising concern about the effectiveness of DIPs as an HIV-1 therapeutic, unless further engineered. Our work further demonstrates that defective proviruses, despite being evolutionary dead ends, are not irrelevant, and prompt future studies to investigate whether RNA and protein expression from defective proviruses contribute to chronic immune activation, exhaustion of anti-HIV immunity, and the development of comorbidities in PWH.

## Materials and Methods

### Study Participants

The study participants were referred by their HIV-1 care providers at the Bartlett Specialty Clinic, Johns Hopkins University. Peripheral blood samples (180mL) were collected at multiple time points. Historical samples were obtained through a longitudinal study at the Bartlett Specialty Clinic. HLA typing was performed by the Johns Hopkins University Immunogenetics Laboratory. Drug concentration levels for P1 were measured by University of Florida Infectious Disease Pharmacokinetics Laboratory.

### Study approval

The Johns Hopkins Institutional Review Board approved this study. All study participants provided written informed consent before enrollment.

### Study of HIV-1 sequences in plasma and CD4^+^ T cells

Whole blood samples were spun at 400g for 10 minutes at 4°C and plasma was removed and spun again for 400g at 10 minutes and frozen at -80°C. Upon thawing, HIV-1 virion-associated RNA was isolated as previously described^106^. RNA was used immediately for reverse transcription with Induro (NEB) according to the manufacturer’s protocol. The cDNA was then used for single genome sequencing as previously described^36^. We initially recovered p6-RT sequences to exclude drug resistance and estimate plasma virus clonality. PCR products obtained with this approach were sequenced by Sanger sequencing (Azenta). CD4^+^ T cells were isolated by magnetic bead-based negative selection (Miltenyi Biotec). Genomic DNA was extracted (QIAamp DNA Mini Kit 51306, Qiagen) from each participant’s CD4^+^ T cells and used for digital PCR (see below) or single genome sequencing. Near-full length HIV-1 proviral sequencing was conducted using a modified version of the FLIP-Seq assay^107^.

### Digital PCR

IPDA and RPP30 normalization was performed as previously described on total CD4^+^ T cells, using the QX200 Digital Droplet^61^. All IPDA experiments included a negative control (nuclease-free water) and a positive control (J-Lat DNA). All RPP30 experiments included a negative control (nuclease-free water) and a positive control (uninfected donor DNA). For P2, we designed custom primers and probes for the *env* region.

Quantification of the participant-specific deletions was conducted by designing custom primers and probes spanning either the 1417 nt or 313 nt (P1), or 270 nt (P2) deletion. For P1, total proviruses was measured using a probe spanning LTR-gag^36^. For P2, we quantified viruses without the 270 nt deletion by designing a custom primer and probe set to bind to the region inside of the deletion (WT-270). To quantify the frequency of proviruses with each respective deletion, we normalized the copies of the deletion targets as a frequency of the total proviruses measured (P1: LTR-gag; P2: WT-270). We also normalized the copies of targets of interests based on cell genome equivalents (calculated by RPP30), as previously described^61^. All dPCR experiments included a negative control (nuclease-free water) and a positive control (diluted amplicon containing deletion of interest). Primer and probes are detailed in Supplemental Table 1. dPCR reactions were run using the QIAcuity One Digital PCR System, with an initial denaturation step of 95°C for 2 minutes followed by 45 cycles of 95°C for 5 seconds and 58°C for 45 seconds. RNA containing each deletion was also quantified (Fig. 3). Virion-associated RNA was isolated as described above. RT-dPCR reactions were run using the QIAcuity One Digital PCR system using the manufacturer’s provided reverse transcriptase and an additional incubation time of 50°C for 40 minutes for reverse transcription before dPCR.

### QVOA

QVOAs from total CD4^+^ T cells were performed as previously described^13^. Supernatants from p24-postive wells were processed as previously described to sequence virion-associated HIV-1 RNA^106^. Primers used to generate amplicons spanning each deletion can be found in Supplementary Table 1.

### Analyses of HIV-1 sequences

Multiple sequence alignments of longitudinal p6-RT or *env* plasma sequences were performed using ClustalW. Root-to-tip distances to a majority consensus sequence for each time point was determined using *p-*distances in MEGA (www.megasoftware.net). Average pairwise distance was determined by averaging the *p-*distance between every sequence pair. A test for panmixia was performed as previously described^108^.

Near-full length amplicons were sequenced were performed by Plasmidsaurus using Oxford Nanopore Technology. Raw FASTQ files were downloaded and either mapped to HXB2 using minimap2^109^ or *de novo* assembled using Flye^110^. Consensus sequences were generated using >75 read coverage. Multiple sequence alignments were performed using MUSCLE. Regions surrounding the recurrent deletions were aligned by hand. Neighbor-joining trees were performed based on a *p*-distance and bootstrap analysis with 100 replicates. Phylogenetic trees were visualized using the *ape* package in R and highlighter plots were generated using a custom script (https://github.com/hariharanviv/highlighterplot).

### Analysis of HIV-1 expression upon T cell activation

Total CD4^+^ T cells were isolated from PBMCs by negative selection. 4M CD4^+^ T cells were plated into a 24-well plate at 2 million/mL in cR10 media (Roswell Park Memorial Institute 1640 Medium with GlutaMAX (ThermoFisher), 10% heat-inactivated fetal bovine serum, 100 U/mL penicillin, and 100 μg/mL streptomycin) with 10 nM dolutegravir, 10 μM tenofovir disoproxil fumarate, 10 μM emtricitabine, and anti-CD3/CD28 antibody-coated magnetic beads (cell-to-bead ratio, 1:1). Cells and culture supernatant were collected after 3 days (P1) or 4 days (P2). Virion-associated RNA was isolated and quantified as described above.

### Testing the impact of deletion signatures on replicative fitness

We used site-directed mutagenesis (NEB) to introduce each major deletion signature into the NL4-3 plasmid, obtained through the NIH HIV Reagent Program, Division of AIDS, NIAID, NIH: Human Immunodeficiency Virus 1 (HIV-1), Strain NL4-3 Infectious Molecular Clone (pNL4-3), ARP-114, contributed by Dr. M. Martin^70^. The primers for the site-directed mutagenesis can be found in Supplementary Table 1. To generate infectious molecular clones, we first seeded 16M HEK293T cells in a T150 flask with 20 mL of Dulbecco’s Modified Eagle Medium with 10% heat-inactivated fetal bovine serum. The next day we transfected the HEK293T cells using lipofectamine 3000, 30 μg plasmid, and 5 μg pAdvantage (Promega). 72 hours after transfection, we collected the supernatant, which was then filtered and concentrated by ultracentrifugation with a 20% sucrose gradient. Virus recovery was measured by p24 ELISA (PerkinElmer).

### Flow cytometry analysis of cells expressing HIV-1 Env

HEK293T cells were transfected with infectious molecular clones carrying the participant-specific deletions (as described above), wildtype plasmid (NL4-3), or an infectious molecular clone with a truncated *env* (NL4-3ΔEnv, a negative control; obtained through the NIH HIV Reagent Program, NIAID, NIH: Human Immunodeficiency Virus Type 1 (HIV-1) Non-Infectious Molecular Clone, pNL4-3ΔEnv, HRP-20281). Transfected HEK293T cells were washed with PBS and dissociated using 2 mL trypsin-EDTA (0.25%) 2 days after transfection. The trypsin-EDTA was quenched by the addition of 8 mL of cD10 followed by washing the cells with PBS. Cells were incubated with 100 μL of yellow-fluorescent reactive dye (Invitrogen; 1:500 dilution in PBS) and incubated for 10 minutes at room temperature in the dark. Next, the viability stain was quenched by the addition of FACS buffer (PBS + 10% fetal bovine serum). Cells were stained with a cocktail of unlabeled primary broadly neutralizing antibodies, 3BNC117 and VRC01 (15μg/mL each) for 1 hour at 37°C. Cells were also stained using the same protocol with broadly neutralizing antibodies (bNAbs) targeting different epitopes (10E8v4, PGT121, PGDM1400, PGT128) at the same concentration. bNAbs were obtained through the NIH HIV Reagent Program, NIAID. After 2 washes, the cells were then stained with APC-labeled secondary antibody against hu-IgG Fc for 30 minutes at 4°C. (100μL of 1:40 dilution; Rat IgG2a, κ; cat 410712 lot B3338999 clone M1310G05). After 2 washes to remove excess antibodies, cells were analyzed using an Intellicyt iQue cytometer. Nonspecific signal was assessed by staining cells with only the secondary antibody.

### Design and production of stable cell lines containing reporter viral genomes with deletion signatures

The plasmid, NL4-3-ΔNef-mTagBFP2 (herein referred to as NL4-3-ΔNef-BFP), was constructed using Gibson assembly to remove the Nef open reading frame from the NL4-3 plasmid and insert mTagBFP2 (Addgene 54572^111^) in the same reading frame. This design links BFP fluorescence with viral infectivity. Variants of this plasmid containing the deletions of interest were generated via site-directed mutagenesis (as described above), resulting in the plasmids NL4-3-ΔNef-BFP Δ1417Δ313 (representing deletions identified in P1) and NL4-3-ΔNef-BFP Δ270 (representing the deletion identified in P2). Due to the impact of these deletions on viral fitness (Fig. 4), a lentiviral system was employed to package these plasmids into lentiviral particles, which were then used to transduce target cells and create a cell line harboring proviruses with the respective deletions. To generate the lentivirus, we transfected HEK293T cells using Lipofectamine 3000 (as described above) with 37.5 μg of the transfer plasmid (either NL4-3-ΔNef-BFP Δ1417Δ313 or NL4-3-ΔNef-BFP Δ270), 10 μg of the packaging plasmid pHelp ^112^, 7.5 μg of pWE, which encodes an HIV-1 CXCR4-tropic envelope, and 5 μg of pAdvantage (Promega). 24 hours post-transfection, the media was removed, the cells were washed with PBS, and fresh media was added (as described above). At 72 hours post-transfection, lentivirus was filtered and concentrated using Lenti-X Concentrator (Takara) according to the manufacturer’s instructions. The lentivirus was resuspended in 1 mL RPMI, aliquoted, and stored at -80°C until used. To generate polyclonal cell lines harboring the defective proviruses, we transduced SupT1 cells (1.5 million cells / 2 mL per well; SupT1 cells engineered to express CCR5; a gift from Dr. James Hoxie^113^) with the previously described lentiviruses packaging defective proviruses. The transduction was carried out by spinoculating SupT1s in the presence of 8 μg/mL polybrene (Sigma-Aldrich), at 1200g for 2 hours at 37°C. The next day, the cells were spun and resuspended in cR10 media containing 10 µM darunavir (MedChem Express) and 10µM enfuvirtide (NIH HIV Reagent Program, NIAID, NIH, HRP-12732, contributed by DAIDS/NIAID) for 3-5 days to eliminate any replication-competent recombinants. Cells with less than 30% transduction efficiency (to reduce the likelihood of multiple lentiviral integrations per cell) as measured by BFP fluorescence were bulk sorted using the MolFlow Legacy and cultured in cR10 medium supplemented with 10% conditioned media.

In vitro *superinfection experiments.* miRFP670 (Addgene 79987^114^) was similarly cloned into the *nef* open reading frame to generate the construct NL4-3-ΔNef-miRFP670 (hereafter referred to as NL4-3-ΔNef-RFP). NL4-3-ΔNef-RFP was transfected following the same protocol, with the exception that 55 μg of NL4-3-ΔNef-RFP and 5 μg of pAdvantage were used during transfection. WT NL4-3-ΔNef-RFP virus was harvested 72 hours post transfection following the same protocol as above.

To test for interference, sorted BFP^+^ cell lines containing a provirus with either deletion signature or mock-transduced (control) were infected with WT NL4-3-ΔNef-RFP virus. 100,000 cells were infected with NL4-3-ΔNef-RFP or mock-infected with cR10 by spinoculation at 1200g for 2 hours at 37°C in a 96-well round-bottom plate in 200 µL cR10 containing 8 μg/mL polybrene. 24 hours after infection, cells were washed with PBS and resuspended in fresh cR10 medium. Daily, 20-50 µL of the cells were analyzed for fluorescent protein reporter expression by flow cytometry (Cytek Northern Lights), with dead cells excluded using propidium iodide staining (BioLegend). On day 3, 50 µL supernatant was removed, spun to eliminate cellular debris, and frozen at -80°C. The titer of day 3 supernatant was determined by using 25 µL of the supernatant to spinoculate 100,000 WT SupT1 cells with the same infection protocol as above. To assess single-round infectivity, 19hrs after infection the supernatant was removed, the cells were washed, and the cells were resuspended in cR10 media containing 10 μM T20 and 5 mM DRV. 48 hours after infection, the frequency of infected cells was measured by flow cytometry (Cytek Northern Lights). The WT titer (Fig. 5c) was determined by normalizing the percentage of cells infected with WT virus (% RFP^+^) or defective virus (% BFP^+^) with condition where SupT1 cells were infected with WT virus only. The fraction of transmitted proviruses (Fig. 5d) was determined by dividing the percentage of cells infected with WT virus (% RFP^+^) or defective virus (% BFP^+^) by the total number of infected cells for each condition (% RFP^+^BFP^-^ + % RFP^-^BFP^+^ + % RFP^+^BFP^+^).

Conditional replication was determined by using the defective cell lines and either mock-infecting them or infecting them with the WT NL4-3-ΔNef-RFP virus. 3 days post infection, the supernatant was removed and used to infect WT SupT1 cells in the same protocol as described above. 24 hours after infection of WT SupT1 cells, the supernatant was removed, and the cells were washed with PBS. Unlike the single-round infectivity experiment, this experiment aimed to assess the replication competence of the viruses, so we resuspended the cells in 200 µL of cR10 medium without antiretrovirals. Daily, 20-50 µL of the cells were analyzed for fluorescent protein reporter expression by flow cytometry (Cytek Northern Lights), with dead cells excluded using propidium iodide staining (BioLegend).

R_0_ quantification by three-color assay was conducted using a modified version of Pitchai, Tanner et al.^50^. First, to generate a target cell line, SupT1-CCR5 were stably transduced with a lentiviral reporter vector, pTY-eGFP (NIH HIV Reagent Program, Division of AIDS, NIAID, NIH: Lentiviral Reporter Vector (pTY-EFeGFP), ARP-4828, contributed by Dr. Lung-Ji Chang) using the same lentivirus protocol and transduction protocol as above in *Design and production of model stable cell lines containing reporter viral genomes with deletion signatures*. Cells with less than 30% transduction efficiency (increasing the probability of only one transgene per cell) were single-cell sorted into cR10 media supplemented with 10% conditioned media and expanded over 28 days. The cell population with the highest percentage of eGFP expression and the highest mean fluorescence intensity (MFI) for surface markers CXCR4 and CCR5 was selected for further experimentation (henceforth referred to as SupT1-eGFP cells). Either WT SupT1s or SupT1-Δ1417Δ313-BFP were infected with WT NL4-3-ΔNef-RFP using the same conditions as above. After 2 days of infection, 30,000 cells were cocultured with 70,000 SupT1-eGFP target cells in 200 µL cR10. Daily, the frequency of infected cells as analyzed as above. R_0_ was calculated by taking the number of secondary infections and dividing it by the number of primary infections. The number of primary WT virus infections was calculated as the frequency of RFP^+^ cells at the time of co-culture (% RFP^+^ × GFP^-^ cells in coculture); the number of secondary infections was calculated by multiplying the frequency of GFP^+^RFP^+^ cells 3 days after co-culture with all uninfected (target) cells (% GFP^+^RFP^+^ × (100,000 – HIV^+^ infected cells at the time of co-culture)). R_0_^DEF^ was calculated similarly, except calculating the frequency of BFP^+^ cells as a measure of infection by the defective virus.

*p24^+^ single cell sorting.* We modified the HIV-Flow assay^77^ to single-cell sort p24^+^ cells into individual wells and lysed them. To ensure that singlets were sorted, we first single sorted live, single cells from an uninfected male donor into 8 μL of lysis buffer (DirectPCR Lysis buffer + 400 μg/mL protease K) and then heated the mixture at 55°C for 60 mins and then 85°C for 15 minutes to inactive the protease K. We added PCR mastermix (Platinum SuperFi II Mastermix) directly to each sorted cell and distributed the mixture across 6 wells, with a final primer concentration of 0.4 μM per well. Primers for amplification of *SRY* are listed in Table S1. We then amplified the product in two rounds of PCR, using the manufacturer’s recommended thermocycling conditions for both rounds and visualized the product by gel electrophoesis. For p24^+^ sorting of ACH-2, P1, and P2 cells, we first activated CD4^+^ cells using 162 nM PMA and 2 μg/mL ionomycin for 24 hours in the presence of 10 nM dolutegravir, 10 μM tenofovir disoproxil fumarate, 10 μM emtricitabine. The next day, extracellular staining was performed using CD3-BV785 (Clone UCHT1), CD8a-APC/Cy7 (Clone RPA-T8), CD4-BV421 (Clone OKT4), CD45RO-FITC (Clone UCHL1), and Live/Dead Fixable Near-IR viability dye (ThermoFisher Scientific cat. L34975). Cells were then fixed and permeabilized with the FOXP3 Buffer

Set (Biolegend cat. 421403), followed by intracellular staining of HIV-1 p24 with clone 28B7 APC (MediMabs cat. MM-0289-APC) and clone KC57 PE (Beckman Coulter cat. 6604667). p24^+^ cells were single sorted into 8 μL of lysis buffer and digested as above. Primers for the amplification of the near-full length genome (FLIP-Seq) and PCR mastermix were added directly to each sorted cell and the mixture was distributed across 6-12 wells, with a final primer concentration of 0.4 μM per well. We then amplified the product in two rounds of PCR, using the manufacturer’s recommended thermocycling conditions for both rounds of PCR. We determined the positive reactions were determined by visualization of amplicons on an agarose gel. Amplicons were sequenced and analyzed as above.

### Data Availability

All HIV-1 sequences are available in NCBI’s GenBank (accession numbers pending).

## Acknowledgments

We deeply thank the study participants for volunteering in this study. We thank Alicia Edwards for administrative support. We thank Natalie McMyn and Joshua Kufera for sharing valuable reagents. We thank Joseph Varriale and Nathan Board for useful discussions leading to this work. This work was also supported by the National Institute on Drug Abuse (U01DA036935) (RDM), the National Institute of Allergy and Infectious Diseases (P01AI169615) (RFS), the Office of the NIH Director and National Institute of Dental & Craniofacial Research (DP5OD031834) (FRS), the Johns Hopkins University CFAR (P30AI094189) (FRS), and by the Howard Hughes Medical Institute (RFS).

## Author contribution

V.H., F.R.S, and R.F.S. conceptualized the study. V.H., J.A.W, F.D., E.J.F., N.P., M.M., and F.R.S. performed experiments. H.Z. conducted cell sorting experiments. A.S., J.L., and S.A.B. enrolled study participants. E.P.S., E.A.G., D.S.B., J.K., and R.D.M. provided clinical care and gathered clinical history. V.H. conducted analyses and generated figures. V.H., J.D.S., R.F.S., and F.R.S. wrote the manuscript with feedback from all authors.

