## Supplemental Figures for "Superinfection with intact HIV-1 results in conditional replication of defective proviruses and nonsuppressible viremia in people living with HIV-1"

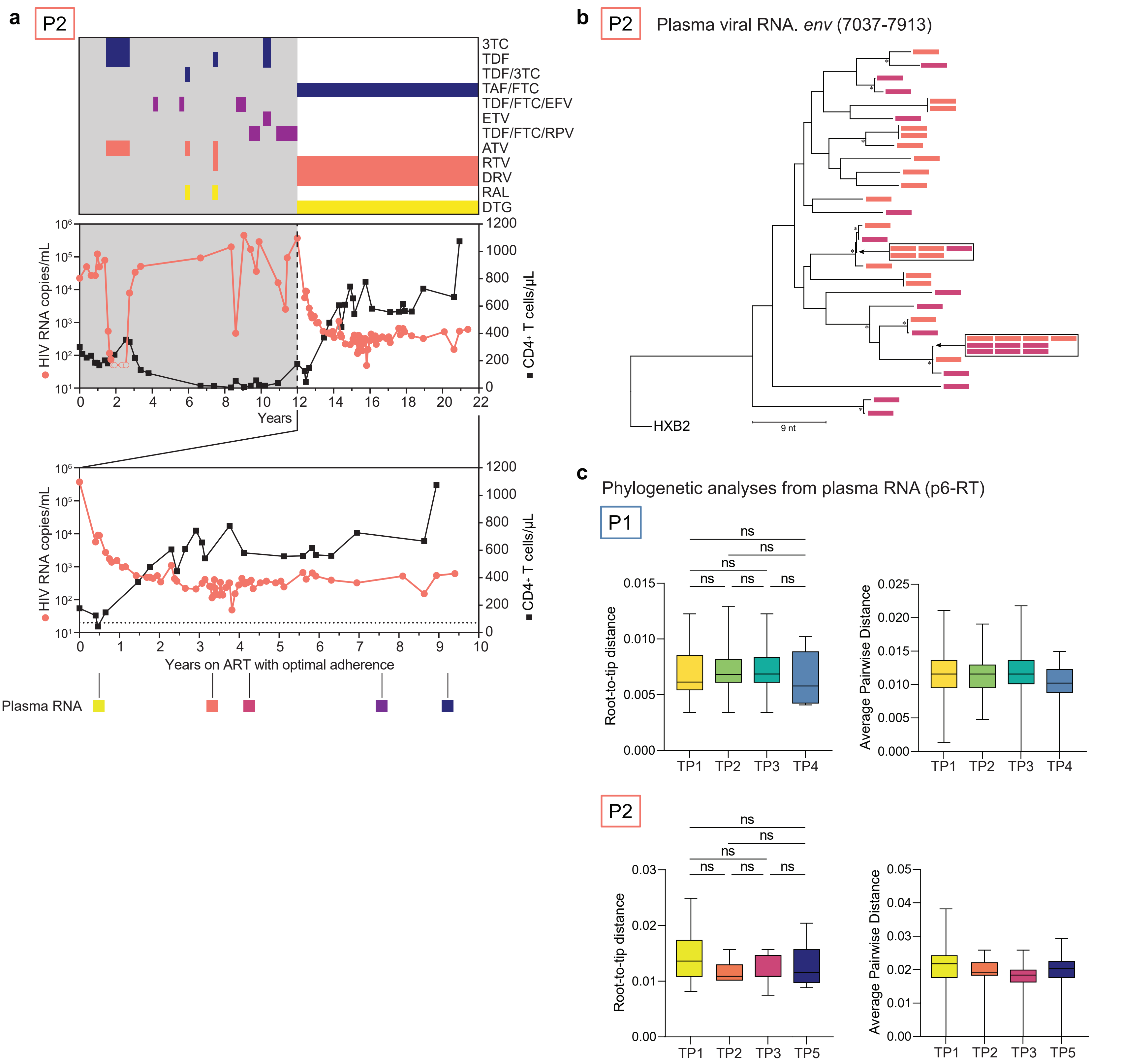


**Supplemental Figure 1. Extended clinical history of P2 and phylogenetic analyses.** (**a**) Extended clinical history for P2. (Top) Prescribed ART regimen for P2. (Middle) Plasma HIV-1 RNA and $CD4^+$ T cell counts for P2. Gray shaded area represents time prior to optimal adherence of ART. (Bottom) Plasma HIV-1 RNA and $CD4^+$ T cell count for P2 for years on ART with optimal adherence (also shown in Fig. 1b). Phylogenetic tree tip labels are color coded according to the plasma collection timepoint in Fig. 1B. (**b**) Neighbor-joining phylogenetic trees of *env* single genome sequences obtained from plasma-associated virions for P2. Tree nodes with bootstrap values above 80 are marked with asterisks. (**c**) Plots of root-to-tip distance and average pairwise distance of P6-RT plasma virus sequences for P1 and P2 showing lack of viral evolution during ART. Box and whisker plots with whiskers representing max to min values. Statistical significance calculated by ordinary one-way ANOVA. n.s.: $P > 0.05$.

3TC: lamivudine; TDF: tenofovir disoproxil fumarate; TDF/3TC: tenofovir disoproxil fumarate/lamivudine; TAD/FTC: tenofovir alafenamide/emtricitabine; TDF/FTC/EFV: tenofovir disoproxil fumarate/emtricitabine/efavirenz; ETV: entecavir; TDF/FTC/RPV: tenofovir disoproxil fumarate/emtricitabine/rilpivirine; ATV: atazanavir; RTV: ritonavir; DRV: darunavir; RAL: raltegravir; DTG: dolutegravir

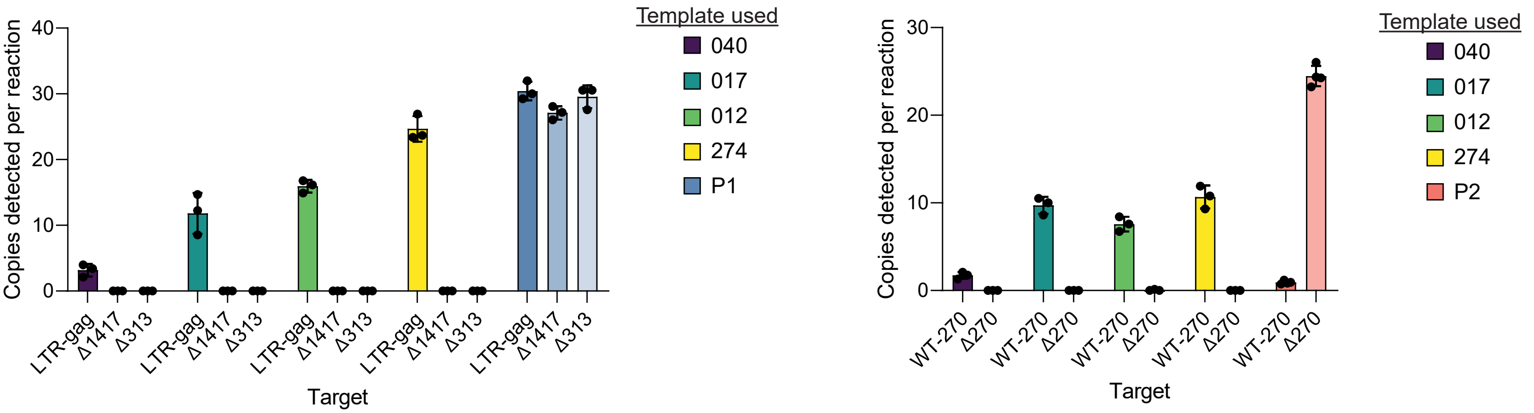


**Supplemental Figure 2. Validation of digital PCR assays.** Results of dPCR reaction using primer and probe sets to specifically amplify deletion regions of interest in 4 PWH on ART (040, 012, 017, and 274), P1, and P2. Horizontal bars represent mean and error bars represent SD.

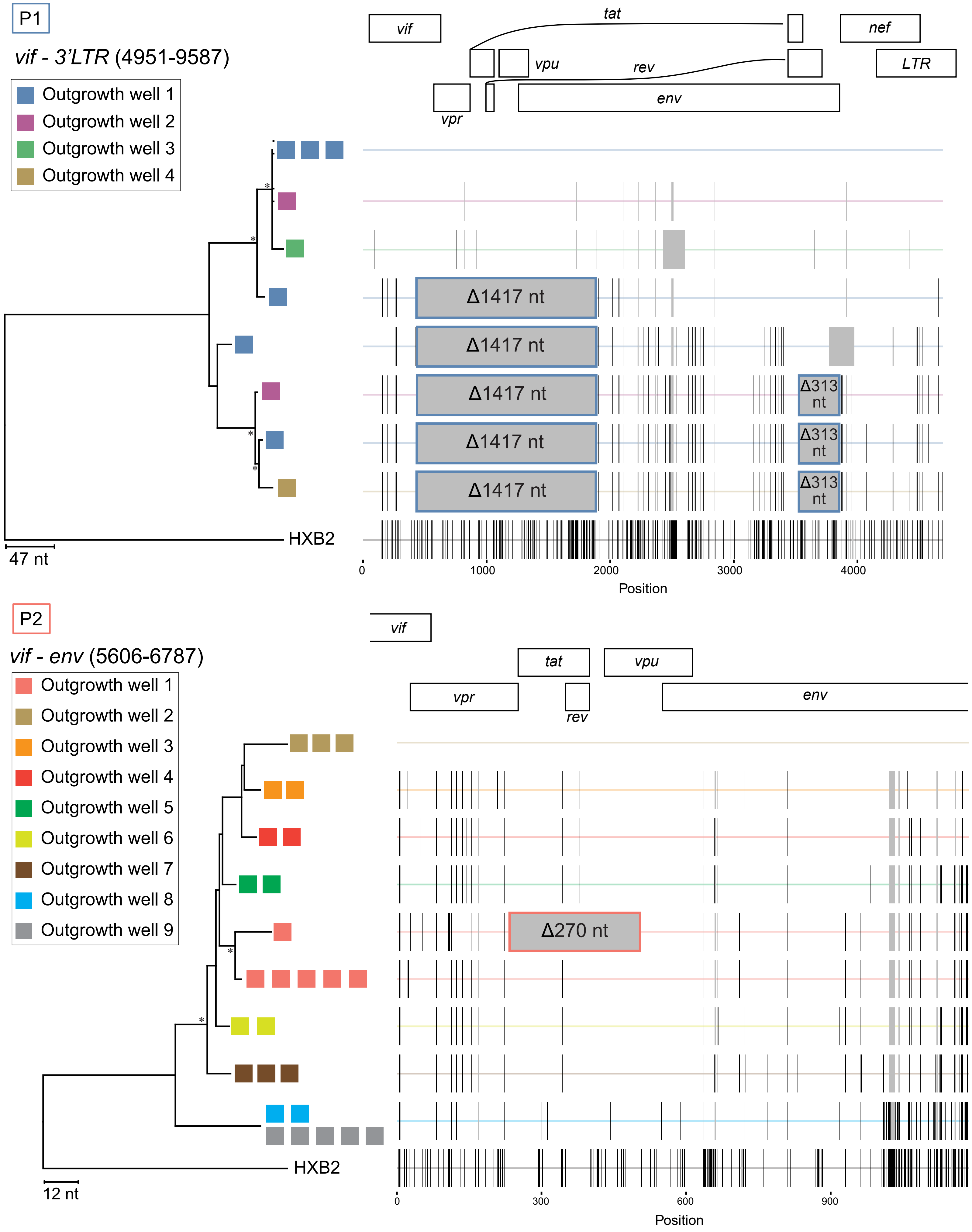


**Supplemental Figure 3. QVOA wells with exponential outgrowth contain proviruses with and without the recurrent deletions.** (**a** and **b**) (Left) Neighbor-joining phylogenetic trees of single genome sequences obtained from outgrowth wells. Colored boxes represent multiple sequences obtained from the same outgrowth well. The sequenced amplicon coordinates refer to the reference genome HXB2. Tree nodes with bootstrap values above 80 are marked with asterisks. (Right) Highlighter plot with black lines representing nucleotide changes compared to the top sequence of each tree. Gray vertical bars represent deletions compared to HXB2. Boxed regions represent recurrent deletion patterns of 1417 nt, 313 nt (P1), and 270 nt (P2).

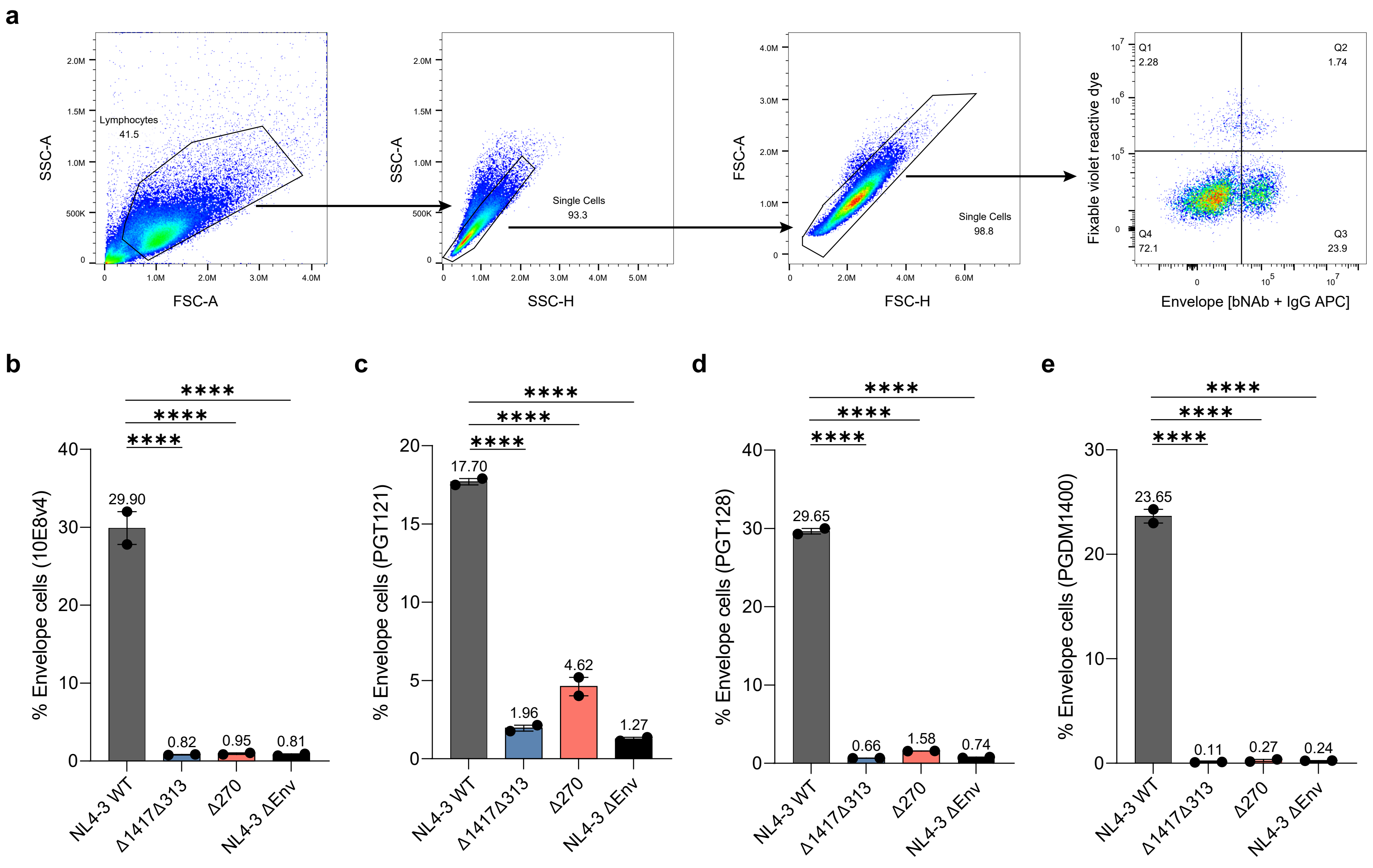


**Supplemental Figure 4. Extended analysis of transfected HEK293T cells.** (**a**) Gating scheme used in flow cytometry to identify transfected cells expressing Env on the cell surface. A representative well of HEK293T cells transfected with NL4-3 WT and stained using a bNAb and secondary α-human IgG is shown here. (**b** - **e**) Surface staining of HIV-1 Env on HEK293T cells 24 hours after transfection with various bNAbs (10E8v4, PGT121, PGT128, and PGDM1400). Horizontal bars represent mean and error bars represent SD. **** $P$ <0.0001

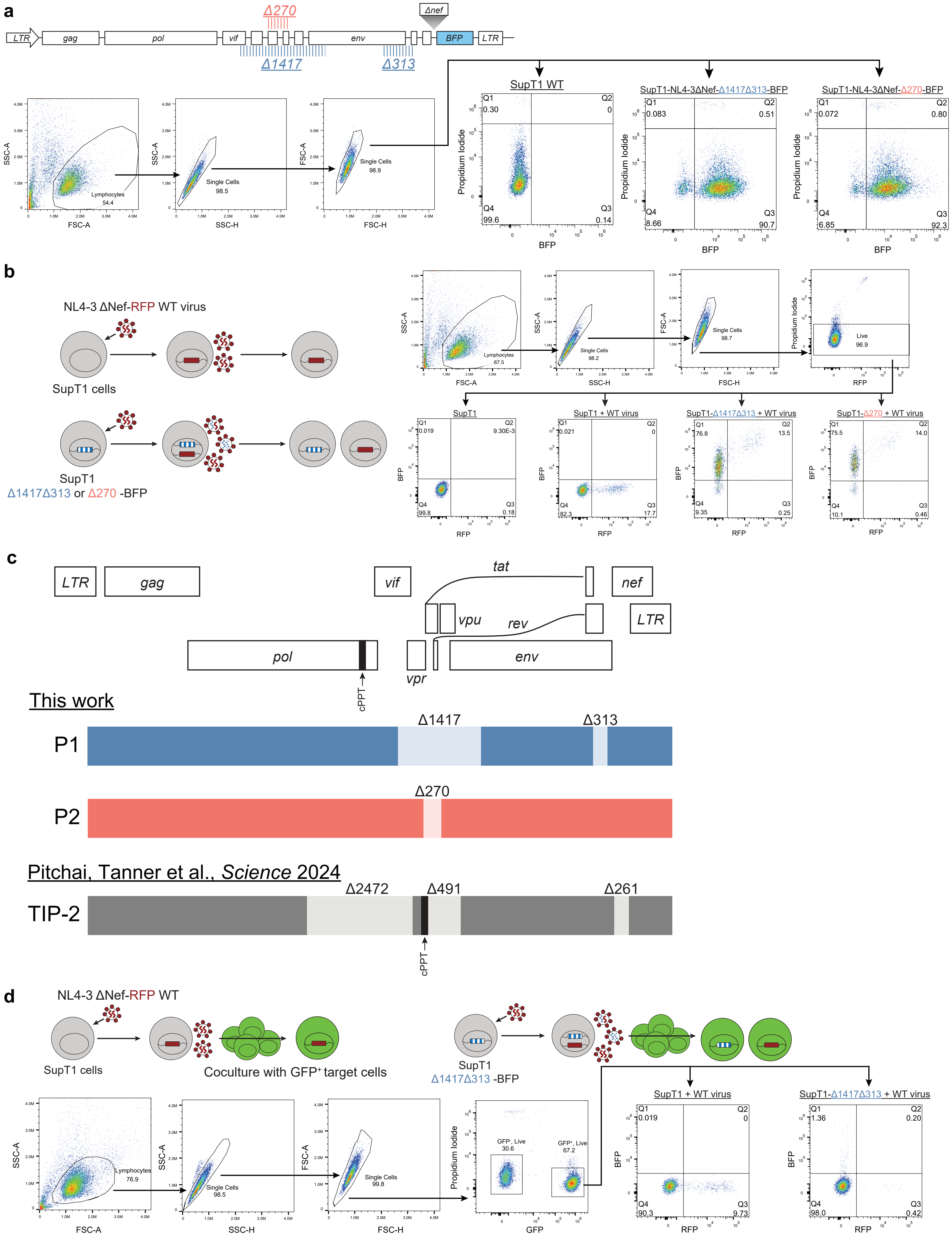


**Supplemental Figure 5. Extended analysis of *in vitro* superinfection experiments.** (**a**) (Top) Construct map of NL4-3-ΔNef-BFP with the deletions marked. (Bottom) Flow cytometry gating of $BFP^+$ sorted cells. (**b**) (Left) Schema of *in vitro* superinfection interference assay. (Right) Representative flow cytometry gating for each condition. (**c**) Comparison of the construct maps showing recurrent deletions identified in P1 and P2 from this study alongside engineered deletions in pNL4-3 from Pitchai, Tanner et al. Light boxes represent deletion regions. The dark black square represents the central polypurine tract (cPPT). (**d**) (Top) Schema of the *in vitro* $R_0$ estimation assay. (Bottom) Representative flow cytometry gating for each condition for the estimation of $R_0$.

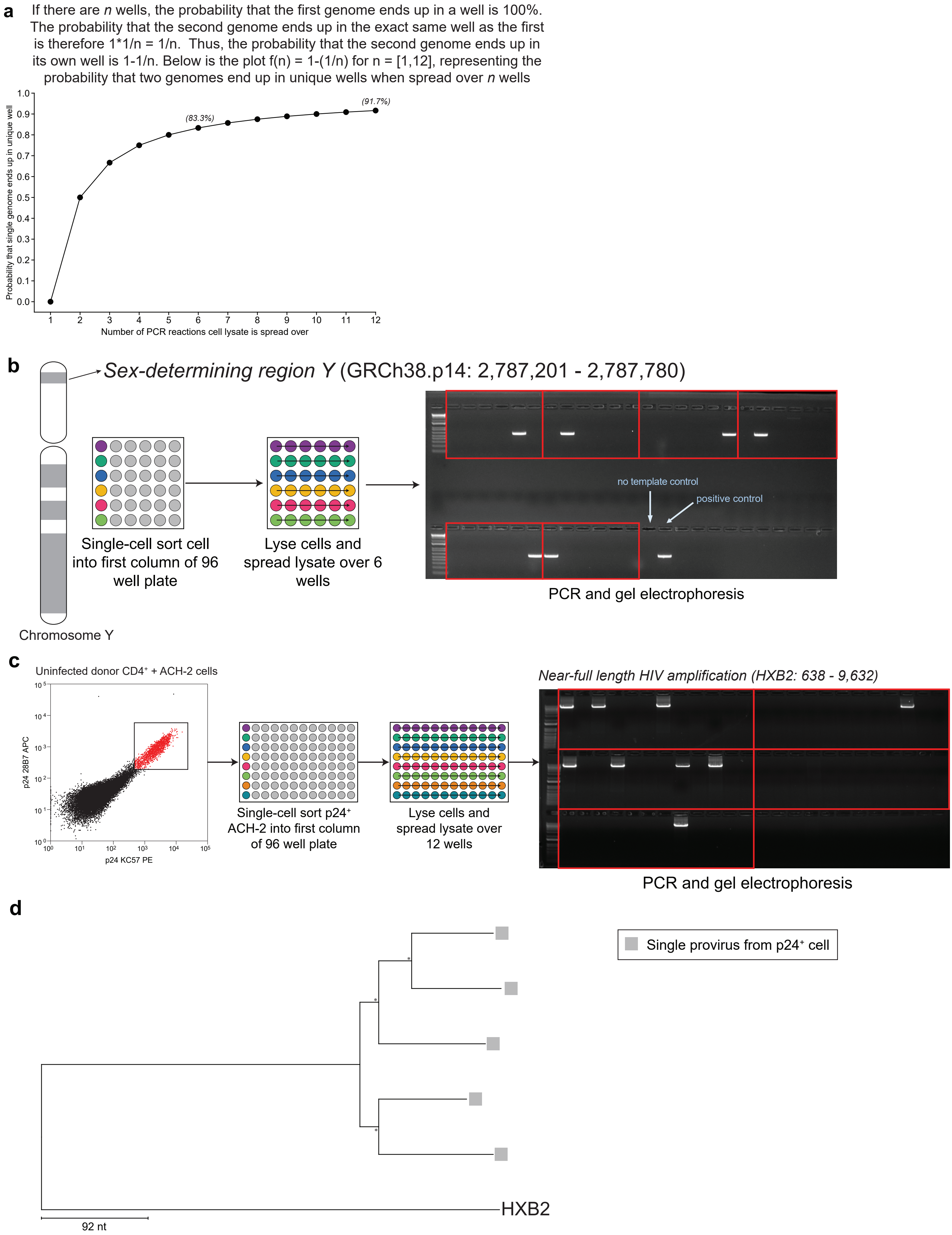


**Supplemental Figure 6. Single cell sorting validation.** (**a**) The probability of separating two genomes, if present, into distinct wells increases as the number of wells over which the cell lysate is distributed grows. (**b**) Live, single, CD4⁺ cells from a male uninfected donor were sorted into lysis buffer and distributed over 6 wells. A region of the *sex-determining region Y* gene was amplified, and the resulting gel shows only one band from each of the 6 sorted cells. Each red box represents the lysate from one cell spread over 6 PCR wells. A no-template control (water only) and positive control (DNA from an uninfected male) are included. (**c**) Uninfected donor CD4+ T cells were mixed with ACH-2 cells, stained for p24, and single-cell sorted. ACH-2 cells were lysed and distributed across 12 wells for near-full-length proviral amplification. The resulting agarose gel shows 12 wells per cell lysate (boxed). (**d**) Neighbor-joining phylogenetic tree from sorted, sequenced p24⁺ cells from a PWH with low-level viremia. Gray boxes indicate a single provirus found in a sorted cell. Nodes with bootstrap values >80 are marked with asterisks.

| Primers used for single genome sequence (SGS) PCR | | | | |
|---|---|---|---|---|
| PCR Target | Primer Name | Primer direction | Primers designation | Primer sequence |
| *Protease-Reverse Transcriptase (P6-RT)* | p6RT_FO | forward | 1st round PCR | GATGACAGCATGTCAGGGAG |
| | p6RT_RO | reverse | 1st round PCR, reverse transcription | CTATYAAGTCTTTTGATGGGTCATAA |
| | p6RT_Fn | forward | 2nd round PCR | GAGTGTTGGCTGAGGCAATGAG |
| | p6RT_Rn | reverse | 2nd round PCR | CAGTTAGTGGTACTATGTCTGTTAGTGCTT |
| Custom *P6-RT* primers for P2 | P2_p6RT_RO | reverse | 1st round PCR, reverse transcription | CTATYAARTCTTTTGATGGGTCATAA |
| | P2_p6RT_Fn | forward | 2nd round PCR | GAGTYTTGGCTGARGCAATGAG |
| | P2_p6RT_Rn | reverse | 2nd round PCR | CTGTTAGAGGKACTACCTCTGTTAGTGCTT |
| *env* | envSGS_fo | forward | 1st round PCR | GCCAGTAGTRTCAACYGAA |
| | envSGS_ro | reverse | 1st round PCR, reverse transcription | GCARATGAGTTTTCYAGAGCA |
| | envSGS_fn | forward | 2nd round PCR | CTGCTAAATGGCAGTCTAGC |
| | envSGS_rn | reverse | 2nd round PCR | TTGCCTGGAGCTGYTTRATGC |
| *vif-3'LTR* | 4813+ | forward | 1st round PCR | GTGCAGGGGAAAGAATAATAGAC |
| | 3LTRi | reverse | 1st round PCR | TCAAGGCAAGCTTTATTGAGGCTTAA |
| | 4930+ | forward | 2nd round PCR | TTTGGAAAGGACCAGCAAAGCT |
| | 3UTRi | reverse | 2nd round PCR | AGGCTTAAGCAGTGGGTTCCCTAG |
| *vif-env* | 5554+ | forward | 1st round PCR | GATAGATGGAACAAGCCCCAGAAG |
| | 6905- | reverse | 1st round PCR | CTTTAGAATCGCAAAACCAGCCG |
| | 5559+ | forward | 2nd round PCR | ATGGAACAAGCCCCAGAAGAC |
| | 6834- | reverse | 2nd round PCR | CCTGTGTGATGCGTGAGGTG |
| Near-full length amplification (FLIP-Seq) | U5-623F | forward | 1st round PCR | AAATCTCTAGCAGTGGCGCCCGAACAG |
| | U5-601R | reverse | 1st round PCR | TGAGGGATCTCTAGTTACCAGAGTC |
| | U5-638F | forward | 2nd round PCR | GCGCCCGAACAGGGACYTGAAARCGAAAG |
| | U5-547R | reverse | 2nd round PCR | GCACTCAAGGCAAGCTTTATTGAGGCTTA |
| *Sex-determining region Y* | SRY_fo | forward | 1st round PCR | AGCAGGGCAAGTAGTCAACG |
| | SRY_ro | reverse | 1st round PCR | TGCAATTCTTCGGCAGCATC |
| | SRY_fn | forward | 2nd round PCR | GGTACTAGGGGGTAGGCTGG |
| | SRY_rn | reverse | 2nd round PCR | CCTTCCGACGAGGTCGATAC |

| Primers and probes used for HIV DNA and RNA quantification | | | | |
|---|---|---|---|---|
| dPCR quantifying Δ1417 | d1417_fwd | forward | digital PCR | CCCTAATCTAGCAGACCAACTG |
| | d1417_rev | reverse | digital PCR | AAGGATATCTTTGGACAGGCTT |
| | d1417_probe | reverse | digital PCR | /56-FAM/TAGTGACTG/ZEN/AGGTGTTACAACTTATCAAAAC/3IABkFQ/ |
| dPCR quantifying Δ313 | d313_fwd | forward | digital PCR | TGAATAGAGTTAGGCAGGGATAC |
| | d313_rev | reverse | digital PCR | CCACCCATCTTATATCAAAGCTC |
| | d313_probe | forward | digital PCR | /HEX/CGACGAAGA/ZEN/AGGTGGAGAGAGAGAAGAA/3IABkFQ/ |
| dPCR quantifying Δ270 | 270_fwd | forward | digital PCR | CAGAATAGGCATTACTCCAAGG |
| | 270_rev | reverse | digital PCR | CTTCTGCTCTTTCACTTATTCTTTC |
| | 270_prb | forward | digital PCR | /56-FAM/AGAAATGGA/ZEN/GCCAATACTAGCAATAGT/3IABkFQ/ |
| dPCR quantifying WT-270 | WT270_prb | forward | digital PCR | /5HEX/TGGAGCCAG/ZEN/TAGATCCTAGACTAGAGC/3IABkFQ/ |
| | WT270_fwd | reverse | digital PCR | TGGGTGTCAACATAGCAGAATAG |
| | WT270_rev | reverse | digital PCR | GCTGACTTCCTGGATGCTT |
| Custom IPDA *env* primers for P2 | P2_IPDA_Env_Fwd | forward | digital PCR | AGTGGTGCAAAGAGAAAAAAGAGC |
| | P2_IPDA_Env_Rev | reverse | digital PCR | GCCTGGCCTGTACCGTCAGC |
| | P2_IPDA_Env_prb | forward | digital PCR | GTTCCTTGGGATCTTGG |
| *LTR-Gag* | LTRgagF | forward | digital PCR | TCTCGACGCAGGACTCG |
| | LTRgagR | reverse | digital PCR | TACTGACGCTCTCGCACC |
| | LTRgagP | reverse | digital PCR | /5TEX615/CTCTCTCCTTCTAGCCTC/3IAbRQSp/ |

| Primers used for site directed mutagenesis | | | | |
|---|---|---|---|---|
| NL4-3-Δ1417 | del1417_F | forward | PCR | GATAAGTTGTAACACCTCAG |
| | del1417_R | reverse | PCR | AAAACAATCAAAATAGTGCAG |
| NL4-3-Δ313 | del313_F | forward | PCR | AGAATAAGACAGGGCTTG |
| | del313_R | reverse | PCR | TCTCTCTCTCCACCTTCTTC |
| NL4-3-Δ270 | del270_fwd | forward | PCR | AATAATAGCAATAGTTGTGTGG |
| | del270_rev | reverse | PCR | GGCTCCATTTCTTGC |

**Supplemental Table S1.** Oligonucleotides used in this study.
